# Targeted syndromic next-generation sequencing panel for simultaneous detection of pathogens associated with bovine reproductive failure

**DOI:** 10.1101/2024.09.10.612295

**Authors:** Dhinesh Periyasamy, Yanyun Huang, Janet E. Hill

## Abstract

Bovine reproductive failure, which includes infertility, abortion, and stillbirth in cattle, leads to significant economic losses for beef and milk producers. Diagnosing the infectious causes of bovine reproductive failure is challenging as there are multiple pathogens associated with it. The traditional stepwise approach to diagnostic testing is time-consuming and can cause significant delays. In this study, we have developed a syndromic next-generation sequencing panel (BovReproSeq), for the simultaneous detection of 17 pathogens (bacteria, virus and protozoa) associated with bovine reproductive failure. This targeted approach involves amplifying multiple pathogen-specific targets using ultra-multiplex PCR, followed by sequencing with the Oxford Nanopore platform and subsequent analysis of the data using a custom bioinformatic pipeline to determine the presence or absence of pathogens. We tested 116 clinical samples and found that BovReproSeq results matched with current diagnostic methods for 93% of the samples, and most of the disagreements occurring in samples with very low pathogen loads (Ct > 35). At the optimal read-count threshold of 10 reads (minimum number of reads to classify the sample as positive), the clinical sensitivity of the assay was approximately 82%, while clinical specificity was 100%. The overall accuracy of the assay was 98.8%. Matthew’s Correlation Coefficient (correlation coefficient of binary classification) was approximately 0.90 and F1 score (harmonic mean of Precision and Recall) was 0.90, indicating excellent overall performance. Our study presents a significant advancement in detecting the infectious agents associated with bovine reproductive failure and the BovReproSeq panel’s ability to detect 17 pathogens makes it a promising tool for veterinary diagnostics.

**Importance:** Bovine reproductive failure causes substantial economic losses to beef and milk producers, and infectious disease contributes significantly to this syndrome. Etiologic diagnosis is complicated since multiple pathogens can be involved and infections with some pathogens are asymptomatic or cause similar clinical signs. A stepwise approach to diagnostic testing is time-consuming and increases the risk of missing the correct diagnosis. BovReproSeq is a next-generation sequencing based diagnostic panel that allows detection of 17 reproductive failure pathogens simultaneously.

## Introduction

Bovine reproductive failure is a collective term used to refer to infertility, abortion, and stillbirth in cattle. It causes substantial economic losses to beef and milk producers, estimated to be approximately $2.8 billion annually in the USA (1). Accurate diagnosis is essential to control and manage this condition. Although non-infectious causes, such as poor nutrition, breeding practices and management, account for most of bovine reproductive failure cases in Western Canada, infectious diseases also contribute to bovine reproductive failure (2). In extreme cases, infectious diseases can lead to abortion storms if not diagnosed correctly. The pathogens associated with this condition also pose a concern to humans as some of them belong to risk group-3 or are zoonotic.

Multiple pathogens are associated with bovine reproductive failure including viruses, bacteria, fungi, and protozoa (Table-1). Infections with some of these pathogens are asymptomatic or cause similar clinical signs and differentiation between these pathogens requires a variety of laboratory tests (Table-1) . There is no recent data available on the diagnostic success rate in Western Canada, however, based on 338 aborted fetuses submitted to Prairie Diagnostic Services Inc. (PDS) in 2010, 52% had no diagnosis, 41% were attributed to an infectious disease, and 7% to a developmental anomaly (2). The success rate for diagnosing bovine reproductive failure is typically low as there is a significant delay between infection and the occurrence of abortion (3). The diagnostic success rate also depends on the availability of appropriate samples (e.g., fresh fetus, placenta, amniotic fluid etc.), and this is not always possible as aborted fetuses are often autolyzed (4). Veterinarians and producers usually focus on testing the most likely cause and if the initial test fails to confirm the suspected cause, subsequent tests are requested. This stepwise approach to diagnostic testing is time-consuming and increases the risk of missing the correct diagnosis as veterinarians and producers are less likely to follow through a diagnostic plan if the initial tests are negative. A syndromic panel approach in which multiple pathogens are tested for simultaneously, would increase the chance of successful diagnosis and significantly reduce the turnaround time for diagnosis (5). Currently, there is no syndromic panel test available in Canada for bovine reproductive pathogens.

PCR is the most preferred method for detection of reproductive failure pathogens due to its high analytical sensitivity and specificity. Even though multiple targets can be detected using PCR, there is a limit to the number of targets that can be included in multiplex assays (6).

Metagenomic next-generation sequencing (mNGS) can overcome this limitation by either sequence-independent amplification of the genetic material in the sample or by directly sequencing the genetic material, thereby providing comprehensive microbial analysis (7). The sensitivity of this method, however, is lower compared to species specific real-time PCR due to the abundance of host genetic material in the sample (8). Sensitivity of mNGS could be improved with higher throughput sequencers and various enrichment methods, such as DNase treatment, rRNA removal, or probe capture; however, this would significantly increase both costs and turnaround times. These limitations can be overcome by combining PCR with high-throughput next-generation sequencing. With this targeted approach, multiple pathogen-specific targets are first amplified in an ultra-multiplex PCR and amplified fragments are then sequenced and analysed (9, 10). While there are several high-throughput sequencing platforms available, the Oxford Nanopore platform offers real-time sequence data and analysis, enabling rapid and timely diagnosis. Multiple samples can be processed in a single sequencing run and diagnostic costs can be significantly reduced. In addition to detecting pathogens of interest, targeted metagenomic sequencing offers the ability to determine strain types (*in silico* serotyping, virulence typing, multi-locus sequence typing (MLST), etc.) and identify variants, which could be useful in differentiating closely related pathogens such as *Campylobacter fetus* subsp. *fetus* and *Campylobacter fetus* subsp. *venerealis* (11). There is also potential to detect emerging or novel pathogens with this approach using random primers along with the targeted primers (12).

In this study, we have developed a syndromic next-generation sequencing panel (BovReproSeq) for simultaneous detection of 17 pathogens associated bovine reproductive failure. The BovReproSeq method was tested on clinical samples, and it successfully detected all 17 pathogens associated with bovine reproductive failure. Despite some limitations in detecting pathogens at very low pathogen loads, the overall performance was excellent with a Matthew’s Correlation Coefficient of 0.90.

## Materials and methods

### Target pathogen selection

Seventeen pathogens including bacteria, protozoa and viruses (both DNA and RNA viruses) were selected for inclusion in the panel (Table 1). Pathogens were selected based on the results of reproductive failure cases that were submitted to Prairie Diagnostic Services since 2014 and additional advice from veterinarians and diagnostic pathologists. Originally 12 pathogens were included and after successful results from initial experiments, five more pathogens were added to the panel bringing the final count to 17 pathogens.

**Table 1:** Pathogens in the BovReproSeq panels and current diagnostic test

| Pathogen | Current diagnostic test |
| --- | --- |
| **Bacteria** |  |
| *Coxiella burnetti* | Real-time PCR |
| *Chlamydia abortus* | Real-time PCR |
| *Ureaplasma diversum* | Real-time PCR |
| *Campylobacter fetus*. subsp. *fetus* | Real-time and conventional PCR |
| *Campylobacter jejuni* | Conventional PCR |
| *Campylobacter fetus*. subsp. *venerealis* | Real-time and conventional PCR |
| *Leptospira* spp. | Immunohistochemistry |
| *Listeria monocytogenes* | Bacterial culture |
| *Trueperella pyogenes* | Bacterial culture |
| *Mycoplasma bovis* | Real-time PCR |
| *Bacillus licheniformis* | Bacterial culture |
| **Protozoa** |  |
| *Neospora caninum* | Immunohistochemistry, serology |
| *Tritrichomonas fetus* | Real-time PCR |
| *Toxoplasma gondii* | Real-time PCR |
| *Sarcocystis* spp. | Immunohistochemistry |
| **Viruses** |  |
| *Bovine alphaherpesvirus 1 (BoHV-1)* | Real-time PCR, virus isolation |
| *Bovine viral diarrhea virus (BVDv)* | Real-time PCR, virus isolation |

### PCR Primer design and *in silico* PCR verification

Multiple PCR assays for each pathogen were identified from published papers. The complete RefSeq genomes for each pathogen were downloaded from NCBI RefSeq database. If complete genomes were not available in RefSeq database, sequences of the target genes or genome contigs of the pathogen were obtained from the NCBI GenBank database*. In silico* PCR was performed in UGENE (version 48) using primer sets from published papers and the downloaded sequences as templates. *In silico* PCR results were evaluated and the primer set that consistently produced the expected amplicon across the highest proportion of representative genomes was chosen for the BovReproSeq panel. A total of 37 primer sets were included based on the *in silico* PCR results, typically two primer pairs targeting each pathogen (Table Sl). For Coccidian parasites (*Neospora caninum*, *Toxoplasma gondii, Sarcocystis* spp.), one specific primer set for each pathogen, and a pan-Coccidia specific primer set were added. An additional seven MLST primer sets were included for *Campylobacter fetus* subspecies differentiation. A partial sequence encoding Enhanced Green Fluorescent Protein (EGFP) was added as an internal amplification control (IC-EGFP) to rule out false negatives due to presence of PCR inhibitors in the clinical samples. Primers for IC-EGFP were designed using primer3 (13). Primers were synthesized by IDT (Integrated DNA Technologies), and 100 μM stocks were prepared for each primer by suspending the lyophilized primer in nuclease free water. Aliquots (5 μl) of each primer stock (100 μM) were pooled together to create a single primer pool.

### Synthetic controls

Synthetic controls were created and used to optimise and validate the targeted next-generation sequencing approach. These controls were generated by flanking random sequences with target-specific primer binding sites. Random sequences with equal nucleotide frequency were generated using the RSAT random sequence web tool (http://rsat.sb-roscoff.fr/random-seq_form.cgi) and then flanked with target-specific primer binding sites (14). The size of each synthetic control was designed to match the actual size of the target (Supplemental File S3). Two to four of these synthetic controls were integrated into a pUCIDT plasmid containing an ampicillin resistance gene, creating a final total of 18 plasmids. *E. coli* DH5-alpha was transformed with these plasmids, and control stocks were stored at -80 °C. Subsequently, plasmids were extracted from overnight *E. coli* cultures in lysogeny broth (LB) containing 100 µg/ml ampicillin using Qiagen QIAprep Spin Miniprep Kit (Qiagen, Cat no. 27104) and plasmid DNA extractions were normalized to 5 ng/μl for further use. Synthetic controls were verified by conventional PCR using both individual and pooled primers. Equimolar amounts of each plasmid were then pooled to make a single pooled synthetic control, ensuring that equal numbers of each synthetic target were in the pooled control. This pooled synthetic control was used as a template for PCR optimisation and as a positive control in clinical sample analysis.

### PCR optimisation with synthetic controls

The aim of this experiment was to determine the optimal multiplex PCR kit, annealing temperature, and primer concentration for the multiplex PCR.Two high-fidelity DNA polymerase kits (Invitrogen Platinum superfi II – Themofisher Cat no. 12361010 – (TF_Plati), and NEB Q5 High-Fidelity 2X Master Mix – NEB Cat no. M0492L (NEB_Q5)) and three annealing temperatures (57 °C, 60 °C, 63 °C) were evaluated. Multiplex PCR was performed with six different combinations of master mixes and annealing temperatures and a primer concentration of 0.015 μM. The synthetic control pool (2 μl) was used as a template. Resulting PCR products (100 ng each sample) from each combination was barcoded with ONT Native barcoding kit (SQK.NBD114.96) and the resulting pooled library was then sequenced on a GridION sequencer using an R10.4 flowcell.

### Reverse-transcription

Following the results from the PCR optimisation experiment, a reverse transcription step was added to the protocol prior to the multiplex PCR with pooled primers to enable the detection of *BVDv* (an RNA virus). Three different reverse transcriptase kits (NEB LunaScript Multiplex One-Step RT-PCR Kit (Cat no. E1555S), NEB LunaScript RT SuperMix Kit (Cat no. E3010L) and NEB LunaScript RT Master Mix Kit Primer-free (Cat no. E3025L) with *BVDv* specific primers) were tested using manufacturer’s instructions with two *BVDv* positive clinical samples.

### Clinical samples

A total of 116 samples were selected from clinical cases submitted to PDS. Each sample was tested for one or more pathogens as requested by the clients using current diagnostic methods (Table 1). This resulted in 152 outcomes: 64 confirmed positive and 88 confirmed negative for the requested tests. Due to the unavailability of positive bovine samples for certain pathogens in the panel, caprine and ovine samples were included, as several abortifacient pathogens are common across cattle, goats and sheep. Various sample types were tested, including placenta, amniotic fluid, vaginal swabs, InPouch media (InPouch TF Bovine, BioMed Diagnostics), semen, and fetal tissues. For pathogens such as Bovine Viral Diarrhea Virus (*BVDv*)*, Mycoplasma bovis,* and *Bovine alphaherpesvirus-1 (BoHV-1)*, positive samples associated with abortions were not available. Therefore, positive samples submitted for respiratory disease investigation were included to ensure representation of these pathogens in the experiment.

Positive clinical samples were not available for *Listeria monocytogenes, Leptospira* spp. (3 different species), *Toxoplasma gondii, Neospora caninum,* and *Sarcocystis* spp. In these cases, mock test samples were created by spiking 10 μl of ATCC culture (*Listeria monocytogenes* (ATCC 7644)*, Leptospira* spp. (3 different species ATCC 23580, ATCC 23606, ATCC 23469), *Toxoplasma gondii* (ATCC 50611), *Neospora caninum* (ATCC 50977), and *Sarcocystis neurona* (ATCC PRA-420)) of the pathogen into 90 μl of placenta matrix (clarified tissue homogenate) that was confirmed negative for BovReproSeq pathogens using the current diagnostic methods in Table 1.

### BovReproSeq workflow

The complete BovReproSeq laboratory workflow includes nucleic acid extraction and reverse transcription, multiplex PCR and library preparation, sequencing and bioinformatic analysis (Figure 1).

**Figure 1:**
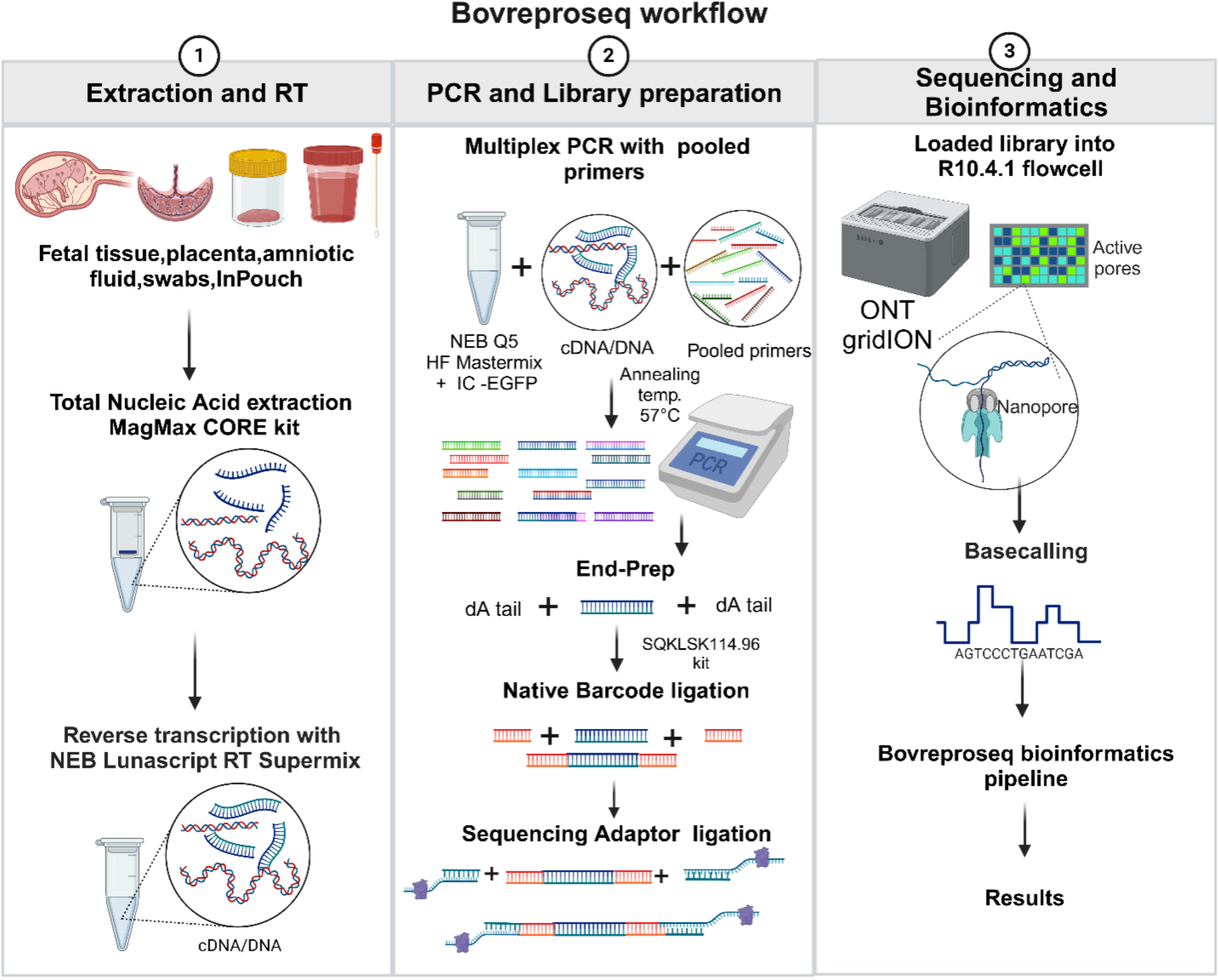
BovReproSeq workflow. 1) Total nucleic acid was extracted from samples using the MagMAX CORE kit, followed by a reverse transcription to enable detection *BVDv*. 2) cDNA/DNA was then used as a template for multiplex PCR with pooled primers and the generated amplicons were then prepared for sequencing using the Oxford Nanopore Ligation Sequencing Kit. 3) The prepared library was loaded onto the R10.4 flowcell and sequenced on the ONT GridION sequencer. The resulting sequencing data were analyzed using a custom bioinformatics pipeline, and a comprehensive results report was generated. (Created with BioRender.com)

#### Total nucleic acid extraction

Applied Biosystems MagMAX CORE Nucleic Acid Purification Kit (Cat no. A32700) was used for extracting total nucleic acid from various sample types, including placenta, amniotic fluid, fetal tissues, and InPouch medium. For placenta and tissue samples, approximately 0.25 g of tissue was added to 1 ml of PBS in a beater tube with glass beads. The tissue was then lysed by beating at 30 Hz for 2 minutes in Qiagen TissueLyser II. The lysate was then centrifuged at 1000 × g for 1 minute, and 100 μl of the clarified lysate (matrix) was used for extraction. In the case of amniotic fluid (200 μl) and InPouch medium (300 μl), the samples were used directly without any pre-processing. Total nucleic acid was extracted from the prepared samples on a KingFisher Flex instrument using the MagMAX CORE kit according to the manufacturer’s instructions.

#### First strand cDNA synthesis

Reverse transcription was carried out with 8 μl of total nucleic acid added to 2 μl of NEB Lunascript RT Supermix (New England Biolabs Cat no. E3010). Nuclease free water was used as a no template control (NTC) and pooled plasmids containing the synthetic controls were used as a positive control. The primers were annealed at 25°C for two minutes, followed by cDNA synthesis at 55°C for 10 minutes and heat inactivation at 95°C for 1 minute. cDNA/DNA from this step was used as a template for the multiplex PCR.

#### Multiplex PCR

The mastermix was prepared by combining 12.5 μl of NEB Q5 Hot Start High-Fidelity 2 × Master Mix (New England Biolabs Cat no. M0492L), 9.5 μl of pooled primers (37 primer sets targeting 17 pathogens, final concentration: 0.5 μM for each primer), 0.9 μl of nuclease-free water, and 0.1 μl of IC-EGFP (150 copies/μl). For each PCR reaction, 2 μl of cDNA/DNA was added to 23 μl of the master mix. The PCR was carried out with the following cycling conditions: initial denaturation at 98°C for 30 seconds, followed by 35 cycles of denaturation at 98°C for 10 seconds, annealing at 57°C for 10 seconds, extension at 72°C for 30 seconds, and a final elongation step at 72°C for 2 minutes. The reaction was then held at 4°C.

#### Magnetic bead clean-up

Agencourt Ampure XP beads (Cat No. Beckman Coulter A63880) (25 μl) were added to each PCR product and mixed thoroughly. The mixture was incubated for 5 minutes at room temperature and then placed on a magnetic rack to separate the beads. Once the solution became clear, it was aspirated and discarded, leaving the pellet. Subsequently, 200 μl of 70% ethanol was added to each sample without disturbing the pellet. The ethanol was aspirated and discarded.

Ethanol washing step was repeated once more. The pellet was allowed to dry for 2 minutes and was then removed from the magnetic rack. The pellet was suspended in 50 μl of nuclease-free water and incubated at room temperature for 5 minutes, followed by magnetic separation. Once clear separation was achieved, the liquid was aspirated and placed into a labeled tube or a 96- well PCR plate. The eluate was then quantified using the Qubit Invitrogen Qubit 4 Fluorometer (Qubit 1x dsDNA HS Assay Kit (Thermofisher, Cat no. Q33231)).

#### Library preparation

The sequencing library was prepared using the ligation sequencing amplicons protocol (Version: NBA_9170_v114_revL_15Sep2022). The cleaned-up PCR product was then normalised to 62.5 ng (200 fmol based on 500 bp fragment size) and taken to end-preparation. The NEBNext Ultra II End repair/dA-tailing Module (NEB, Cat no. E7546) was used to add dA-tails to the ends of the amplicons, followed by magnetic bead clean up and elution with 12 μl of nuclease free water. Subsequently, unique barcodes from the Native Barcoding Kit 96 V14 (ONT, Cat No. SQKNBD114.96) were ligated to the end-prepped amplicons using NEB Blunt/TA Ligase Master Mix (NEB, Cat no. M0367). After barcoding, samples with unique barcodes were pooled together and cleaned up using magnetic beads. Sequencing adapters were then ligated to the pooled samples using the NEBNext Quick Ligation Module (NEB, Cat no. E6056), followed by magnetic bead clean up. Finally, the resulting library was quantified using Invitrogen Qubit 4 Fluorometer with the (Qubit 1x dsDNA HS Assay Kit (Thermofisher, Cat no. Q33231)).

#### Sequencing

The prepared library (20 ng (∼ 65 fmol)) was loaded onto a ONT R10.4.1 flowcell and sequenced on a GridION platform for a minimum duration of 15 hours. Super-accurate basecalling option in MinKNOW was used for basecalling. Only reads with barcodes on both ends were used and barcodes were trimmed while sequencing using the trim barcodes option in MinKNOW software (Versions used: 23.04.5, 23.04.6, 23.07.5, 23.07.12, 23.11.3)

#### Bioinformatics analysis

A Nextflow bioinformatics pipeline was developed for the analysis of sequencing data and interpretation of results (Figure 2). The path to a directory containing subdirectories with fastq files and the path to a Kraken2 Plus-PF-8 database (k2_pluspf_08gb_20230605) (15) were used as the inputs for the pipeline. Individual fastq files were merged followed by quality filtering (q20) for each sample using nanoq (version- 0.10.0) (16). The resulting merged file was then mapped to a reference sequence containing the amplicon sequences using minimap2 (version 2.25)(17). SAMtools (version-1.17) was used to generate read statistics and split the bam file with aligned reads into amplicon-specific bam files (18). Primer trimming was performed on both ends of the sequences using ampliconclip option in SAMtools (version-1.17). Consensus sequences were generated on the split bam files with a minimum mapping quality of 30 (19) and a base quality of 10 using SAMtools (version-1.17) (18). These generated consensus sequences were then polished using medaka (version-1.11.3) (20) and compared against a custom Blast database containing the BovReproSeq amplicon sequences using ABRicate (version-1.0.1) (21, 22) to determine the presence or absence of the pathogen and internal amplification control (IC- EGFP). Concurrently, taxonomic classification was performed on the raw reads using Kraken2 (version-2.1.3) (15), to identify the unmapped reads and to support the results from ABRicate (version-1.0.1). The results from kraken2 (version-2.1.3) were visualized as an interactive pie chart using krona (version-2.7.1) (23). Output data from these analyses are collated into a HTML report. All the quality metrics from SAMtools and Nanoplot (version-1.42.0) were captured and visualised using multiQC (version-1.14) (24, 25). The pipeline, along with detailed information, is accessible on GitHub at https://github.com/dhineshp565/BovReproSeq.

**Figure 2:**
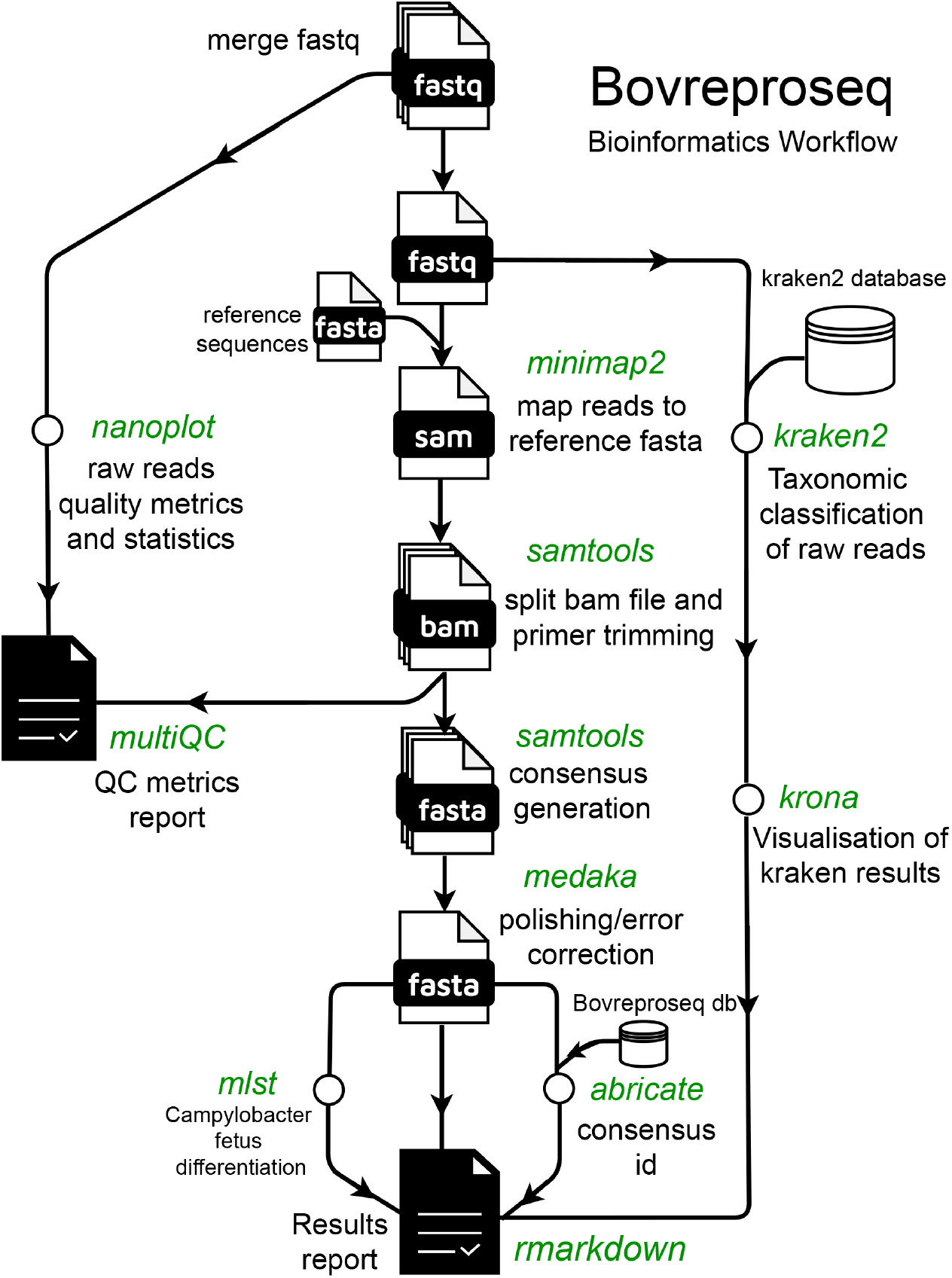
BovReproSeq Bioinformatics workflow. Multiple FASTQ files generated for each sample were merged into one and renamed. The merged file was then mapped against a reference using minimap2. The generated SAM file was subsequently used to generate consensus and read statistics using SAMtools. The consensus sequence was further analyzed using MLST and ABRicate to determine the type and identity of each consensus. In parallel, merged reads were taxonomically classified using kraken2. Finally, all the results were stitched together to create a comprehensive results report.

### Real-time PCR

Real-time PCR was performed on clinical samples to verify results from BovReproSeq. Total nucleic acid was extracted using Applied Biosystems MagMAX CORE Nucleic Acid Purification Kit (Cat no. A32700) and used as the template for real-time PCR. For DNA-based assays, real-time PCR was performed using Applied Biosystems TaqMan Fast Advanced Master Mix (Cat No: 4444965), with specific primers and TaqMan probes tailored to each pathogen.

Detection of Bovine Viral Diarrhea Virus (*BVDv*) was carried out using the ThermoScientific VetMAX-Gold BVDV PI Detection Kit (Cat no: 4413938).

### Evaluation of read-count threshold and overall performance

Read-count thresholds represent the minimum number of reads required to classify a sample as positive or negative. To determine whether a read-count threshold of 10 reads is optimal for classifying a sample as positive for a target, sequence data from clinical samples were analyzed using the BovReproSeq pipeline with 12 different thresholds (ranging from 1 read to 100 reads). The results from each threshold were then compared to the available real-time PCR results. For each read-count threshold, a contingency matrix (2x2 table) was created to check the agreement of results between the two methods. Due to the imbalance between positive and negative results, a precision-recall curve was used instead of a receiver operator characteristic curve to determine the optimal read-count threshold (26, 27). Performance metrics, including sensitivity, specificity, accuracy, and precision were calculated for each read-count threshold using the data from 2 x 2 contingency table. Precision-recall curves were plotted to evaluate the performance of the BovReproSeq method across different thresholds. The F1 score (harmonic mean of precision and recall) was used to determine best read-count threshold. An F1 score of 1 represents perfect precision and recall, indicating optimal model performance, and a F1 score of 0 occurs when either precision or recall (or both) is zero, indicating poor model performance (28). Matthew’s correlation coefficient (MCC) was calculated to determine the overall performance of the assay. MCC is a specialized application of Pearson’s correlation coefficient, designed for imbalanced binary classification scenarios (28, 29). It incorporates all four categories of the contingency matrix (True Positives, False Positives, True Negatives, and False Negatives) to provide a comprehensive score of overall performance (28, 29). The contingency table, Precision-Recall curves, F1 score and MCC were generated using custom Python scripts with the following libraries: pandas for data manipulation (30), scikit-learn for performance metric calculations (31), and matplotlib for visualization (32).

## Results

### PCR optimization with synthetic controls

The BovReproSeq method was developed and optimized using synthetic controls before testing the approach on clinical samples. Pooled plasmids containing synthetic control sequences for 12 pathogens and one internal amplification control (IC-EGFP) were used as a template to test two different multiplex PCR kits (NEB_Q5 and TF_Plati), and three different annealing temperatures. The total mapped reads obtained from each amplification protocol were as follows: NEB_Q5_57°C: 439,971, TF_Plati_57°C: 409,795, NEB_Q5_60°C: 470,324, TF_Plati_60°C: 414,916, NEB_Q5_63°C: 458,083, and TF_Plati_63°C: 486,733. The sequencing depth of each amplicon varied significantly in each combination (Figure 3). The NEB_Q5 with an annealing temperature of 57°C yielded a higher read depth for most of the amplicons (17 out of 29) and a balanced performance, with reads evenly distributed across different amplicons compared to other treatments, which showed skewed distributions favoring specific amplicons (Figure 3). As a result, the NEB Q5 High-Fidelity 2x Master Mix and an annealing temperature of 57°C were used in subsequent experiments. The read depths for *Campylobacter fetus* targets *glyA, aspA*, and *Ureaplasma diversum 16S rRNA* were particularly low (<500 reads) compared to other targets in all six combinations of annealing temperatures and polymerases. To address this, primer concentration of each primer was increased to 0.5 μM and new primer pool was made. The multiplex PCR was then repeated with NEB Q5 High-Fidelity 2x Master Mix with an annealing temperature of 57°C and new pooled primers (0.5 μM), which resulted in a significant increase in the number of reads mapping to *Campylobacter fetus glyA* (40 reads to 1047 reads), *aspA* (137 reads to 2670 reads), and *Ureaplasma diversum 16S rRNA* (44 reads to 1214 reads) (Figure 4).

**Figure 3:**
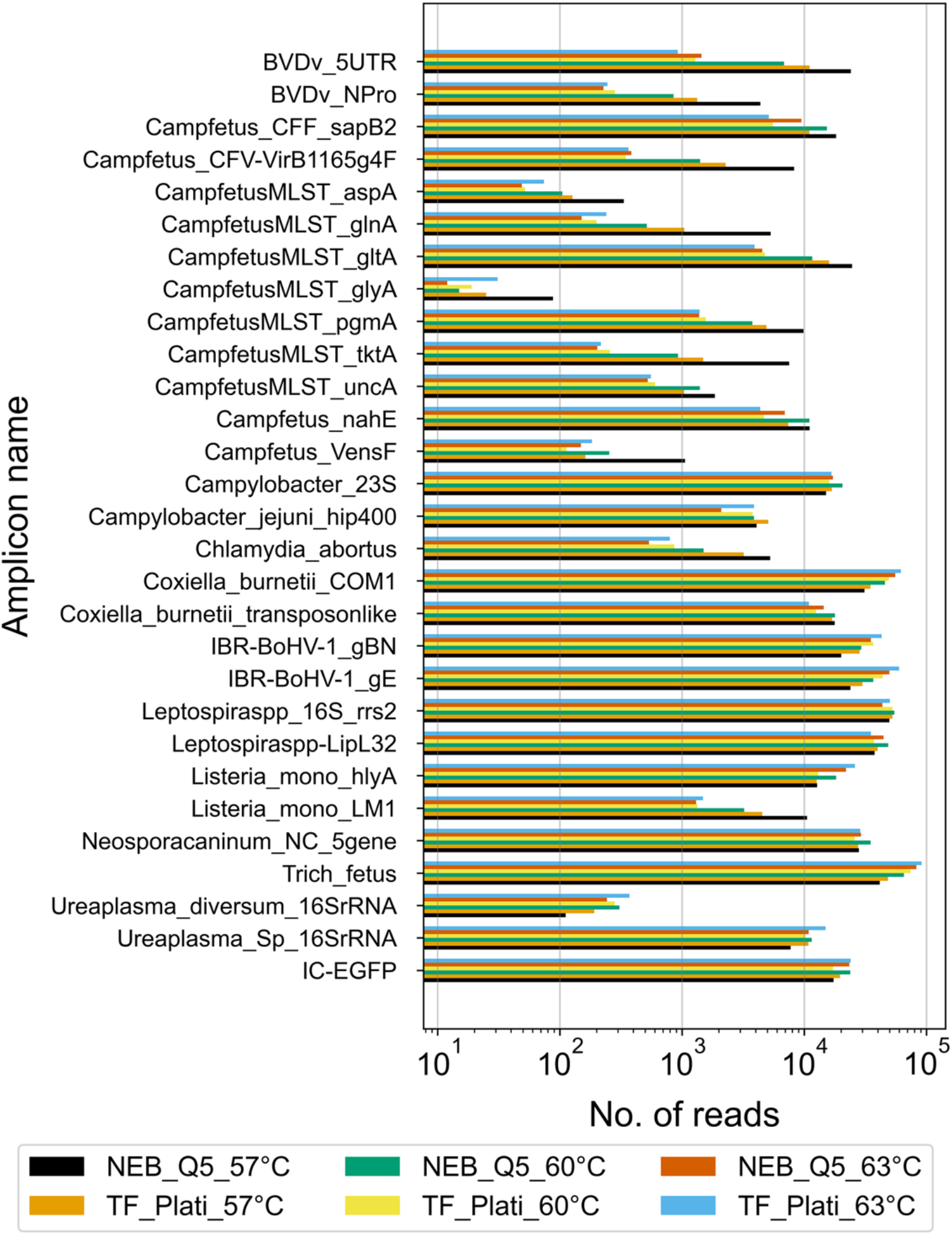
Comparison of sequencing depth of amplicons produced with NEB Q5 High-Fidelity 2x Master Mix and TF Invitrogen Platinum superfi ii polymerase, annealing temperatures of 57°C, 60°C and 63°C, and primer concentration of 0.015 μM. NEB Q5 High-Fidelity 2x Master Mix at 57°C consistently produced higher read depths across most amplicons. This experiment was performed without the reverse- transcription step and includes results only from synthetic controls for 12 pathogens (29 synthetic targets).

**Figure 4:**
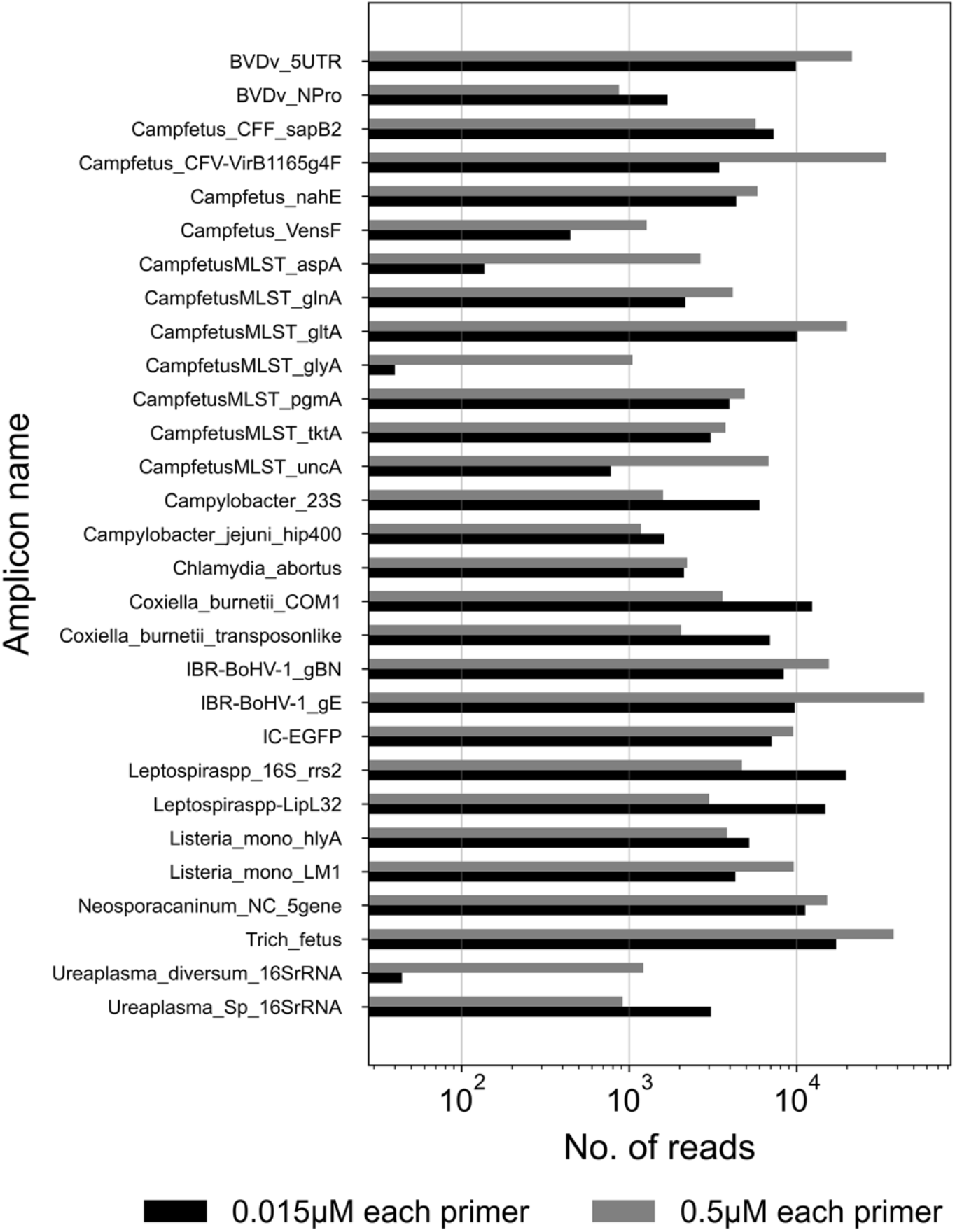
Comparison of sequencing depth for amplicons produced with two different primer concentrations (0.015 μM and 0.5 μM) using NEB Q5 High-Fidelity 2x Master Mix and an annealing temperature of 57°C. The higher primer concentration of 0.5 μM consistently yielded greater read depths across most amplicons compared to the lower concentration of 0.015 μM. This experiment was performed without the reverse-transcription step and includes results only from synthetic controls for 12 pathogens (29 synthetic targets).

As a result, primer concentration of 0.5 μM was used in subsequent experiments. Following these optimizations, five additional pathogens (*Toxoplasma gondii*, *Sarcocystis* spp., *Bacillus licheniformis, Trueperella pyogenes and Mycoplasma bovis)* were added to the panel, and new synthetic controls were created for these pathogens. The updated pooled synthetic control was tested, and all 37 synthetic targets were detected.

A reverse transcriptase step was then added to the multiplex PCR step to enable the detection of *BVDv* (RNA virus), and three different reverse transcriptase kits were tested on two *BVDv* positive clinical samples. Gel electrophoresis results from the NEB LunaScript Multiplex One-Step RT-PCR Kit (Cat no. E1555S) indicated a significant amount of short non-specific amplifications, so the PCR product from this kit was not sequenced. PCR products from the NEB LunaScript RT SuperMix Kit (Cat no. E3010L) and the NEB LunaScript RT Master Mix Kit (Primer-free, Cat no. E3025L) with *BVDv*-specific primers were sequenced. The NEB LunaScript RT Master Mix Kit (Primer-free) yielded 10,598 *BVDv*-specific reads in sample-1 and 7,385 *BVDv*-specific reads in sample-2. In contrast, the NEB LunaScript RT SuperMix Kit with random hexamers produced 1,170 *BVDv*-specific reads in sample-1 and 2,567 *BVDv*- specific reads in sample-2 (Table S4). A control experiment without the RT step showed no detection of *BVDv* in either sample. Despite the lower read depth for *BVDv*, the NEB LunaScript RT SuperMix Kit (Cat no. E3010L) was selected because it successfully amplified *BVDv* RNA in clinical samples and allows for the potential detection of additional pathogens not included in the panel, due to the presence of random hexamers in the mix.

### IC-EGFP titration and repeatability

An internal amplification control was included in the BovReproSeq panel to rule out false negatives caused by inhibiting factors such as humic acid and urea in clinical samples. The amount of IC-EGFP to be spiked into each reaction was determined experimentally, as excessive amounts could interfere with the detection of targets of interest, while insufficient amounts might go undetected.

To determine the optimal quantity of IC-EGFP to be spiked into each sample, a titration experiment was conducted using two concentrations of IC-EGFP: 150 copies/reaction and 15 copies/reaction. A mock *Neospora caninum* positive sample was prepared by spiking 10 μl of *Neospora caninum* ATCC 50845 culture into 90 μl of placenta matrix (clarified tissue homogenate) confirmed negative by current diagnostic methods (Table 1). This mock sample was serially diluted (10-fold dilution) using placenta matrix (clarified tissue homogenate) as the diluent to maintain consistent matrix composition across dilutions. Total nucleic acid (DNA/RNA) was extracted from the dilutions, and BovReproSeq PCR was performed in triplicate for each of the two IC-EGFP concentrations. For each concentration, a sequencing library was prepared and sequenced on separate flow cells*. Neospora caninum* real-time PCR was performed in triplicate to determine the Ct value of each dilution.

Both pan-Coccidia and *Neospora caninum*-pNC-5 gene targets were consistently detected in all replicates and dilutions up to a Ct value of 32 (10^-5^ dilution) in both experiments demonstrating the repeatability of the assay (Figure 5A, 5B). However, for dilutions with Ct values above 32, detection of both *Neospora caninum* targets was inconsistent, occurring only in one or two of the replicates. As the concentration of *Neospora caninum* decreased in the dilutions, the number of IC-EGFP reads increased in both experiments. The IC-EGFP reads were at their lowest when the number of reads and concentration of *Neospora caninum* were at their highest (Ct 18.25).

**Figure 5:**
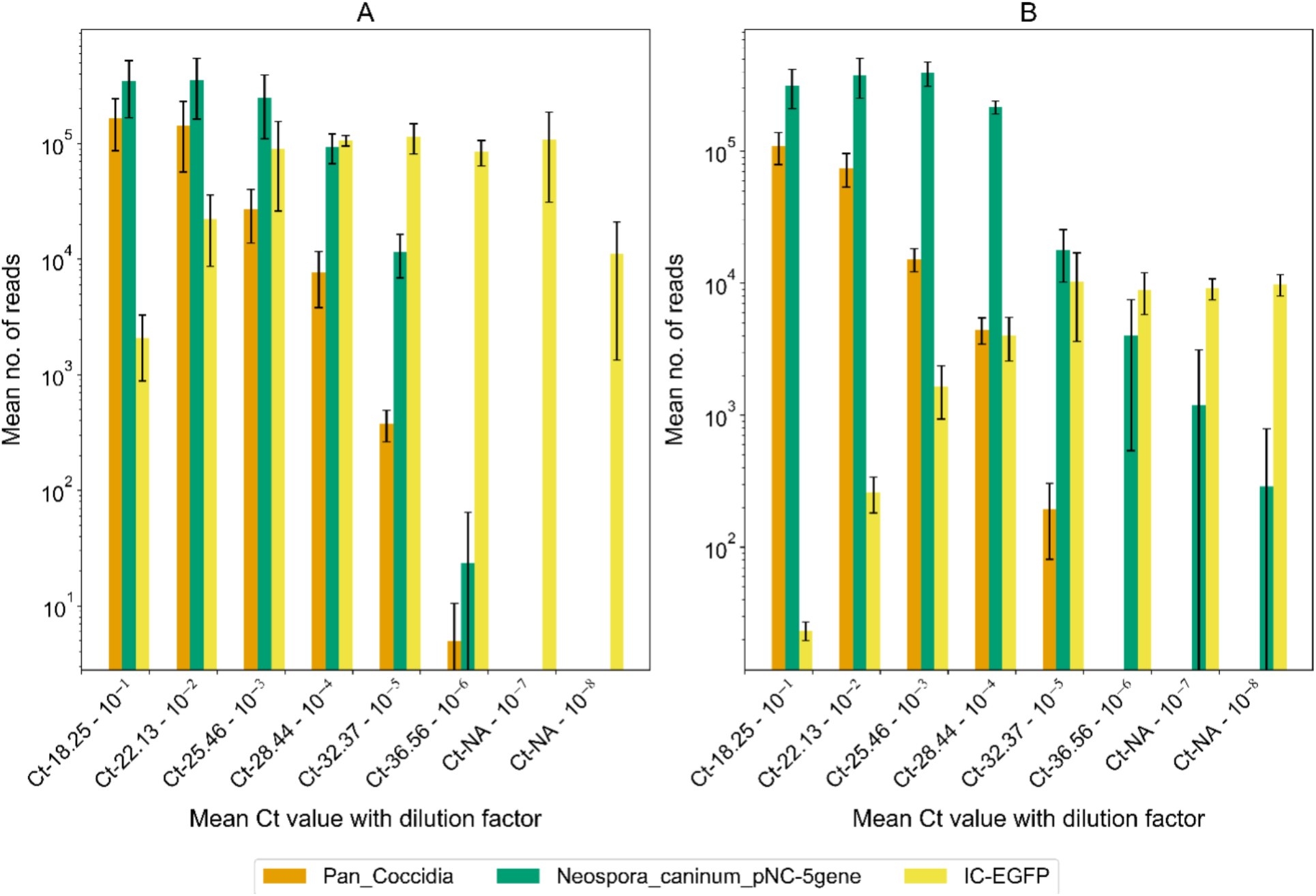
Comparison of mapped reads in serially diluted mock *Neospora caninum* positive samples spiked with 150 copies (A) and 15 copies (B) of IC-EGFP per reaction. Each dilution was tested in triplicate. The x-axis shows mean Ct values from real-time PCR of serial dilutions of the mock sample while y-axis represent the mean number of reads generated for each amplicon from BovReproSeq. “NA” on X-axis indicates that *Neospora caninum* was not detected using real-time PCR. Error bars represent the standard deviation of read counts across triplicates. *Neospora caninum* was consistently detected in all replicates up to Ct of 32, with inconsistent detection in subsequent dilutions. 15 copies of IC-EGFP per reaction yielded higher read depth for *Neospora caninum* in higher dilutions than 150 copies per reaction.

Specifically, in the 15 copies/reaction experiment, there were 23 IC-EGFP reads out of 42,275 total reads, while in the 150 copies/reaction experiment, there were 2,081 IC-EGFP reads out of 51,175 total reads. In both experiments, IC-EGFP reads stabilized around a dilution of 10^-5^. The experiment with 15 copies of IC-EGFP per reaction demonstrated better read depth for *Neospora caninum*, particularly in dilutions with lower pathogen load (Ct 36.56). At this dilution, the 15 copies/reaction experiment yielded 4,021 *Neospora caninum* reads out of 12,932 total reads, compared to only 21 *Neospora caninum* reads out of 85,252 total reads in the 150 copies/reaction experiment. Moreover, *Neospora caninum* was detected by BovReproSeq in dilutions 10^-7^ and 10^-8^ which were not detected with real-time PCR. Although these results suggest that the BovReproSeq method is more sensitive than real-time PCR, it is important to note that these experiments used samples spiked with live organisms. The sensitivity may vary in actual clinical samples, where factors such as sample type, sample matrix and presence of inhibitors could influence detection limits. Based on these results, 15 copies of IC-EGFP per reaction was selected as the optimal concentration for subsequent experiments, as it provided a better balance between internal amplification control and pathogen detection.

### Specificity evaluation

A total of 29 clinical samples, both positive and negative for BovReproSeq targets, along with no-template controls (NTC) and pooled BovReproSeq synthetic positive controls, were tested to determine the specificity of the assay. Results from the BovReproSeq method matched current diagnostic methods for all samples (Table S5). However, most of the samples showed a high abundance of reads mapping to the *Campylobacter jejuni* amplicon (cjejuni23s) (Table S5).

Manual analysis of the consensus sequence using BLAST revealed that these reads did not correspond to *Campylobacter jejuni*. Instead, they matched other *Campylobacter* species and *Campylobacter*-like organisms. Similar non-specificity issues were observed with the Tpyo- cpn60 target, as reads mapping to this target did not match *Trueperella pyogenes* reference sequences and resulted in poor consensus sequences. Additionally, in *Mycoplasma bovis* (Mbovis) positive clinical samples, the Mbovis gltX target was consistently undetected, and the gspA target was detected with only a very low number of reads. Due to the non-specificity of cjejuni23s, Tpyo-cpn60, and the poor performance of both Mbovis targets, these primers were replaced with more specific targets to improve the overall assay performance. Synthetic controls were developed for the newly added targets, and a new set of pooled controls was made (Table Sl). This updated pooled synthetic control and the clinical samples were then tested with BovReproSeq method. All 37 synthetic controls including the replacements were detected from a single PCR reaction (Figure 6).

**Figure 6:**
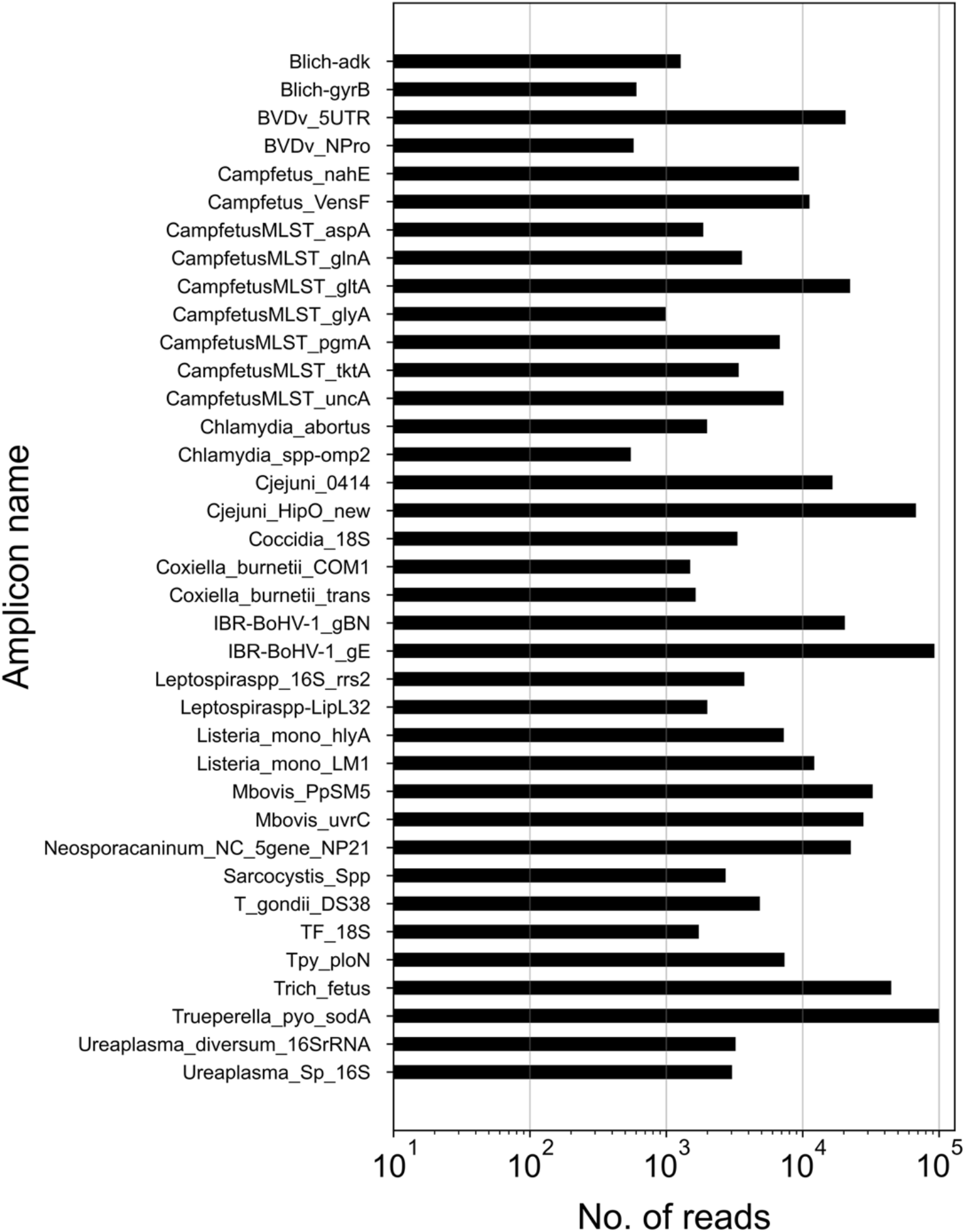
Comparison of read counts for each synthetic control from a single PCR reaction (including reverse transcription step), where the final version of the pooled control was used as a template. All 37 synthetic targets were detected from a single PCR reaction.

### Performance on clinical samples

To evaluate the performance of the BovReproSeq method against current diagnostic methods (Table 1), 116 clinical samples with existing results (for tests requested by veterinarians and pathologists) were tested using the BovReproSeq approach. BovReproSeq PCR was performed and PCR amplicons from these samples were pooled into four libraries (∼30 samples per library), each library loaded onto a ONT R10.4.1 flow cell, and sequenced. The data was analyzed using the BovReproSeq bioinformatics pipeline. The number of raw reads per sample varied significantly, ranging from 6,636 to 642,695 reads, with a median of 110,509 reads. The median total bases generated per sample was 105.5 Mb, and the median raw read quality score was 19.6. Positive controls had reads and consensus sequences mapping to all 37 synthetic controls, while negative controls (NTCs) were negative for all pathogens. Two samples lacked IC-EGFP reads, indicating potential PCR inhibition, and were omitted from further analysis. The results from the BovReproSeq method matched the existing results from current methods for approximately 93% of the samples (139/150, Table 2). The kappa value of 0.85 indicated almost perfect agreement between the two methods, however, there were 11 false negatives. The Ct values from the current diagnostic PCRs for positive samples ranged from 12 to 37, with all false negatives having Ct values between 33 and 37, which is consistent with low pathogen load (Figure 7).

**Figure 7:**
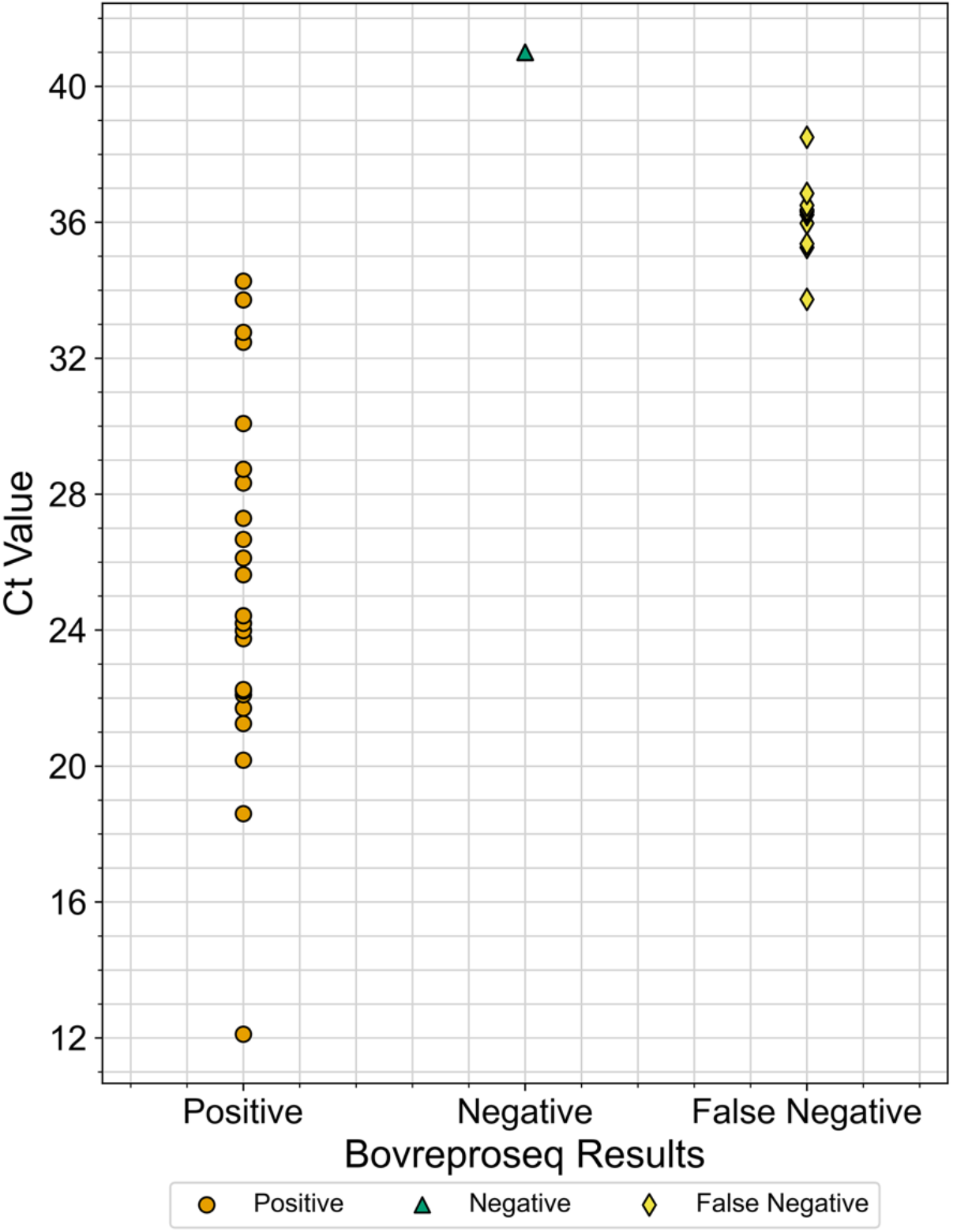
Real-time PCR Ct values of the samples and the corresponding results from the BovReproSeq method. Ct values for the positive samples ranged from 12 to 36, while all the false negatives had a Ct value close to 35 or above indicating that these samples had a very low pathogen load. Additionally, some of these false negative samples had a high abundance of reads mapping to pathogens such as *Ureaplasma diversum* and *Trueperella pyogenes*.

**Table 2:** Comparison of results between current laboratory methods and BovReproSeq. It includes only the results for which specific testing was requested.

|  |  | BovReproSeq results |  |  |
| --- | --- | --- | --- | --- |
|  |  | Positive | Negative | Total |
| Current lab methods | Positive | 52 | 11 | 63 |
|  | Negative | 0 | 87 | 87 |
|  | Total | 52 | 98 | 150 |

BovReproSeq method successfully differentiated *Campylobacter fetus* subspecies in 6 cases and identified the MLST type in five of them (Table S2). Additionally, BovReproSeq identified *Pestivirus tauri (BVDv*-type 2) in two samples among those tested for *BVDv* (Table S2).

Notably, in 43 samples, additional pathogens were detected using the BovReproSeq method that were not part of the initial testing request (Table S2). In most cases, these additional pathogens were either *Ureaplasma diversum* or *Trueperella pyogenes*. However, in seven cases where the initial test requested was negative, abortifacient pathogens such as *Coxiella burnetii*, *Campylobacter jejuni*, or *Campylobacter fetus* were identified (Table 3). Subsequent real-time PCR results confirmed the presence of these additional pathogens.

**Table 3:** Comparison of results from initial test request (current methods) and BovReproSeq method.

| Sample ID | Tests requested by veterinarians (Initial test request) | Results from current methods | Results BovReproSeq |
| --- | --- | --- | --- |
| 6135-2 | *Chlamydia abortus*, *Coxiella burnetii* | Negative | *Campylobacter jejuni* |
| 6156-2 | *Chlamydia abortus*, *Coxiella burnetii* | Negative | *Campylobacter jejuni* |
| 6157-2 | *Chlamydia abortus* | Negative | *Campylobacter jejuni* |
| 6783-3 | *BVDv*, *Campylobacter fetus* subsp. *venerealis*, *Coxiella burnetii*, *BoHV-1*, *Tritrichomonas fetus* | Negative | *Trueperella pyogenes* |
| 1581-2 | *Coxiella burnetii* | Negative | *Chlamydia abortus*, *Trueperella pyogenes* |
| 2760-2 | *Coxiella burnetii* | Negative | *Campylobacter fetus* subsp. *fetus* |
| 8292-1 | *Ureaplasma diversum* | Negative | *Coxiella burnetii* |
| 4122-2 | *Bovine alphaherpesvirus 1 (BoHV-1)* | Positive | *BoHV-1*, *Mycoplasma bovis*, *Coxiella burnetii*, *Trueperella pyogenes* |
| 4123-2 | *Bovine alphaherpesvirus 1 (BoHV-1)* | Positive | *BoHV-1*, *Ureaplasma diversum*, *Mycoplasma bovis*, *Trueperella pyogenes*, *Pestivirus tauri (BVDv type-2)* |

The presence of additional pathogens and the detection of pathogens in samples where the initial test request was negative indicated that to comprehensively evaluate the BovReproSeq method, each sample had to be tested for all 17 pathogens using conventional methods. This would provide a true comparison between the current methods and the BovReproSeq method. This was not possible as some of the samples used were archived nucleic acid samples collected over a long period, limiting our ability to apply methods other than PCR. Real-time PCR assays were available for only 10 BovReproSeq pathogens (*BVDv, Campylobacter jejuni, Campylobacter fetus* subsp. *venerealis, Neospora caninum, Toxoplasma gondii, Coxiella burnetii, Chlamydia abortus, Tritrichomonas fetus, Ureaplasma diversum, Mycoplasma bovis*). Therefore, real-time PCR was performed on samples (109 samples) that had enough nucleic acid to detect the presence of these 10 pathogens, providing 1090 results. Results from each PCR were collated into a single results table (1090 PCR results). The compiled real-time PCR results for 10 pathogens across 109 clinical samples showed 1016 negative and 74 positive outcomes. These real-time PCR results were then compared to the results from the BovReproSeq method. The results from the BovReproSeq method matched the current methods for 98.8% of the samples (1077/1090, Table 4). There were 13 false negative results. All false negative results occurred in samples with Ct values between 33 and 37 except for one sample with a Ct value of 29 for *BVDv.* In this case, four other pathogens (*BoHV-1, Mycoplasma bovis, Coxiella burnetii* and *Trueperella pyogenes*) were detected in high abundance, and the pathologist had reported lesions characteristic of *BoHV-1*. The kappa value of 0.90 indicated almost perfect agreement between the two methods. Due to the imbalance between positive and negative results, the kappa value may not accurately reflect the performance of the assay (*33*). Therefore, Matthew’s Correlation Coefficient (MCC) was calculated. MCC value of approximately 0.90 indicated excellent overall performance of the assay.

**Table 4:** Comparison of results between Real-time PCR assay of 10 pathogens and BovReproSeq.

|  |  | BovReproSeq results |  |  |
| --- | --- | --- | --- | --- |
|  |  | Positive | Negative | Total |
| Real-time PCR results | Positive | 61 | 13 | 74 |
|  | Negative | 0 | 1016 | 1016 |
|  | Total | 61 | 1029 | 1090 |

### Evaluation of read-count threshold and assay performance

Precision (positive predictive value) and recall (sensitivity) were calculated for read-count thresholds from 1 to 100 reads as described in the Methods, and a precision-recall curve was plotted (Figure 8). At the lowest threshold of one read, precision was at its lowest around 48%, while sensitivity peaked at 95%. Increasing the read-count threshold to five reads resulted in a substantial improvement in precision (86%) but a marked reduction in sensitivity (85%).

**Figure 8:**
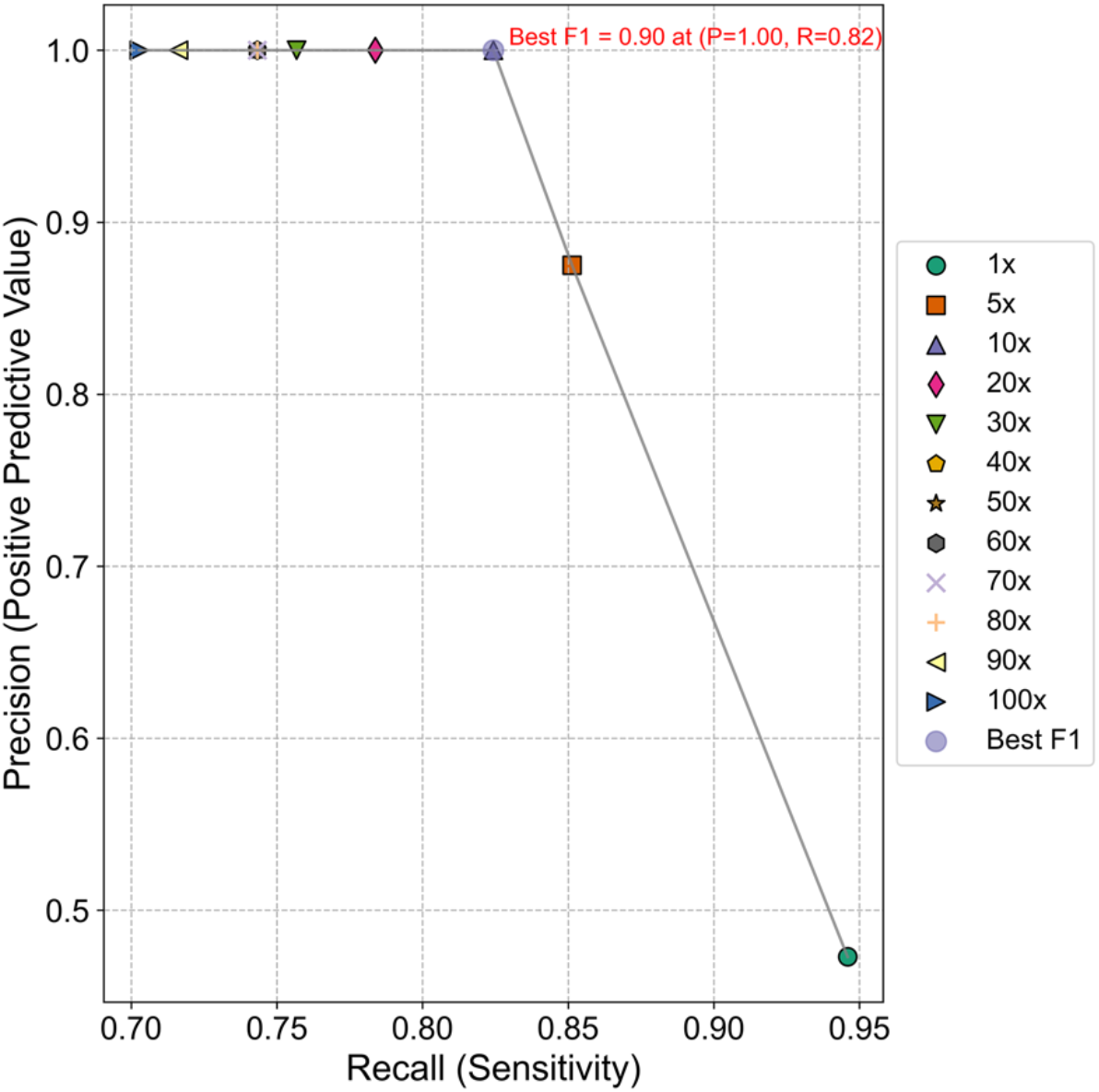
Precision-Recall curve. The graph shows the trade-off between precision (positive predictive value) and recall (sensitivity) at various read-count thresholds. The grey line represents the precision-recall curve, with data points corresponding to different read-count threshold (1x to 100x). The optimal performance point (10x) is highlighted, showing the best F1 score of 0.90 with a precision of 1.00 and recall of 0.82.

Precision reached 100% at a threshold of 10 reads and remained consistent across remaining thresholds, but sensitivity declined with increasing read-count thresholds and was lowest (70%) at a threshold of 100 reads. The threshold of 10 reads yielded the optimal balance between precision and recall with the maximum F1 score of 0.90. At the read-count threshold of 10 reads, the sensitivity of the assay was approximately 82%, while specificity was 100%. The positive predictive value was 100%, and the negative predictive value was 98.7%. The overall accuracy of the assay was 98.8%. Based on these results, a read count threshold of 10 was determined to be optimal for the BovReproSeq method.

## Discussion

Our study demonstrates the development and validation of a syndromic next-generation sequencing panel (BovReproSeq) designed to simultaneously detect 17 pathogens associated with bovine reproductive failure. Initial testing with a pooled synthetic control successfully identified all 37 synthetic targets in the products of a single PCR reaction, demonstrating the multiplexing ability of the assay. The results from BovReproSeq testing of 116 clinical samples matched with existing results from current diagnostic methods for 93% of samples. Additionally, the BovReproSeq method detected pathogens not originally requested for testing, indicating possible mixed infections (Table S2). This ability of the BovReproSeq to identify additional pathogens highlights its advantage over current single-plex diagnostic methods by providing broader detection without iterative testing.

In addition to detection of 17 pathogens, BovReproSeq method enables the typing of *BVDv* (*Pestivirus bovis* (*BVDv* type-1) and *Pestivirus tauri (BVDv* type-2)) based on differences in 5′- UTR sequences of *Pestivirus*. The 5′-UTR sequences could also be used to detect *Pestivirus braziliense* and *Border disease virus* in sheep and goats. With *Campylobacter fetus*, in addition to detection and differentiation of subspecies, we are also able to identify the sequence type using MLST (Table S2), which aids in epidemiological studies. Since this assay involves random hexamers and multiple primers, there is potential to detect pathogens not included in the panel, further broadening the scope of the assay. The entire BovReproSeq workflow (Figure 1) can be completed within 48 hours of receiving the sample and can be cost-effective if multiple samples are processed within a single flow cell.

The BovReproSeq method has some limitations, particularly in samples with high Ct values (close to or above 35) indicating that very low pathogen concentrations in the sample may not be detected (Figure 7). This finding is consistent with a similar study in which a targeted next-generation sequencing method was not able to detect vector-borne pathogens in canine samples with Ct values greater than 35 (10). It is worth noting that PCR artifacts and cross-contamination can sometimes lead to amplification at Ct values >35, and samples with Ct values >35 are often considered “suspect” rather than “positive” in diagnostic laboratories. Moreover, pathogens like *Coxiella burnetii* and *Chlamydia abortus* are known to occur in healthy animals in low numbers and their presence in low quantities without associated lesions is generally not considered as the cause of the abortion (34). In most infectious abortion cases, high pathogen loads with characteristic lesions or signs are typical. Thus, the failure of BovReproSeq to detect very low abundance pathogens may not be a clinically significant limitation. Notably, we had only one sample (Sample ID - 4122-2) with a Ct value of 29 for *BVDv* where BovReproSeq failed to detect the virus. In this case, four other pathogens in the BovReproSeq panel (BoHV-1, *Mycoplasma bovis, Coxiella burnetii* and *Trueperella pyogenes*) were detected in high abundance, and the pathologist had reported lesions characteristic of BoHV-1 (Table 3).

Similarly, we frequently detected pathogens such as *Ureaplasma diversum* and *Trueperella pyogenes*, and in a couple of cases, their abundance had a negative impact on the detection of the causative pathogen. This suggests that multiple pathogens in high abundance in a sample may impede the detection of pathogens in low abundance. This limitation could be overcome by removing primers for *Ureaplasma diversum* and *Trueperella pyogenes* from the main pool and placing them in a separate PCR reaction, which could reduce their impact on the detection of other targeted pathogens.

The analytical sensitivity of our BovReproSeq method is lower compared to PCR assays. The sensitivity of detection of specific pathogens in highly multiplexed assays are significantly impacted by the amount and number of other targets that are in the sample (35). This limitation could be addressed by reducing the number of samples per flow cell, extending sequencing time to improve the sequencing depth of low abundance targets and by using high-throughput sequencers such as ONT PromethION, Illumina MiSeq or NovaSeq. The one-pot multiplex PCR could also be split into multi-pot PCR with smaller primer pools to improve sensitivity, and the number of PCR cycles could be increased from 35 to 40 to increase the number of copies of the amplicons. Regarding primers and PCR target selection, selecting specific genes of pathogens rather than conserved housekeeping genes was more successful, as the primers targeting conserved genes (*T. pyogenes* cpn60 and *C. jejuni* 23S rRNA gene) were more likely to amplify sequences from related organisms present in the sample and obscure the targeted pathogens of interest. This observation should be factored into any future additions to or modification of the panel.

Similar studies have been published recently, but our research presents a few methodological differences and advancements (Anis et al., 2018; Kattoor et al., 2022). Anis et al. developed a targeted NGS method for the simultaneous detection of 43 common bovine and small-ruminant pathogens. Their primer design was a collaboration with the White Glove Team (Ion Torrent; Thermofisher Scientific) using Ion Ampliseq Designer (Ion Torrent; Thermofisher Scientific, Waltham, MA). Kattoor et al. focused on the comprehensive detection of 17 vector-borne pathogens in canines. The primers in their study were designed with Thermofisher Scientific’s AgriSeq Bioinformatics team, and the primer sets are now proprietary items of Thermofisher (Kattoor et al., 2022). Both studies used Ion Torrent Suite and Geneious software for bioinformatics and neither includes an internal amplification control. In our study, we have used previously published primers and further evaluated them based on *in silico* PCR results.

Additionally, in the BovReproSeq bioinformatics pipeline, we have used open-source tools and packaged them as one Nextflow pipeline that produces a readable results report and the pipeline can be scaled up to run on any platform (36). Our pipeline is currently available on GitHub, and the primer information is included in the supplementary section (Table S1) to ensure open access to our methods and facilitate reproducibility.

The previously published bovine and canine pathogen sequencing panels used Thermofisher Ion Torrent technology with IonChef for sequencing multiplex PCR products. This second-generation sequencing technology has limitations including short reads (<400 bp), low throughput and poor accuracy in homopolymer regions. BovReproSeq uses Oxford Nanopore technology (ONT), a long-read approach that enables sequencing of longer gene segments than the Ion Torrent platform and allows for real-time sequencing and analysis, resulting in faster turnaround times. Additionally, BovReproSeq uses only full-length reads to generate the consensus sequences resulting in highly accurate results. The use of long reads allows us to perform MLST typing, as most MLST fragments are around 600 bp, which would be challenging with Ion Torrent technology. Although the raw read accuracy with ONT was known to be lower compared to other platforms, the raw read quality has significantly improved with the newest version of the R10 nanopore and the associated kit 14 chemistry (37, 38). It is also important to note that the raw read accuracy of long homopolymer regions is still poor with ONT; however, this is not an issue with BovReproSeq, as the amplicons are relatively short and do not contain long homopolymer regions. In our BovReproSeq pipeline, we exclusively used raw reads with a quality score of 20 or above (>99% accuracy) to ensure that only highly accurate reads were used for downstream analysis.

The incorporation of an internal amplification control in the BovReproSeq assay allows the rule out of false negatives due to PCR inhibition. The amount to be spiked into each reaction was determined experimentally and to ensure repeatability, we performed this assay in triplicates on serial dilutions with consistent results between the replicates (Figure 5A, 5B). Additionally, the pooled synthetic positive control for the assay allows the detection of positive control contamination since the sequences of the positive control amplicons are unrelated to the actual biological targets. In contrast, using the actual pathogen sequence as a positive control could complicate interpretation. If contamination occurs from the positive control, a positive result could be attributed to either the contamination of the assay or the genuine presence of the pathogen. Therefore, using a synthetic control that shares the same primer binding sites but has different sequences in between will help identify contamination while also making sure that PCR is working correctly. Finally, we analysed the sequence data generated from 109 clinical samples using 12 different read-count thresholds to determine the optimal threshold and the overall performance of assay, significantly enhancing the reliability and robustness of our results.

## Conclusion

In conclusion, our study presents a significant improvement to diagnosis of infectious causes of Bovine reproductive failure. The ability of the assay to detect 17 pathogens despite some limitations, makes BovReproSeq panel a promising tool for veterinary diagnostics. This approach could be applied to other syndromic diseases and could contribute to broader improvements in animal health diagnostics. However, as with all nucleic acid-based methods, the detection of a pathogen does not necessarily mean it is the cause of the disease and results from the BovReproSeq method must be carefully interpreted with pathological findings and case history.

## Data availability

All synthetic control sequences, reference amplicon sequences, and the custom ABRicate database with Accession numbers for each sequence, along with the Nextflow pipeline, are available in the GitHub repository at https://github.com/dhineshp565/BovReproSeq.

## Acknowledgements

We thank Saskatchewan Agriculture Development Fund (ADF) and the Beef Cattle Research Council (BCRC) for their financial support. We also extend our gratitude to the PDS Molecular Diagnostics section for providing the samples and assisting with follow-up real-time PCR, and Anju Tumber for her help in procuring reagents.

