## Supplementary File S3 for "Targeted syndromic next-generation sequencing panel for simultaneous detection of pathogens associated with bovine reproductive failure"

### Bovreproseq synthetic positive control sequences

Synthetic controls were created and used to optimise and validate the targeted next-generation sequencing approach before testing it on clinical samples. Random sequences with equal nucleotide frequency were generated using the RSAT random sequence web tool (http://rsat.sb-roscoff.fr/random-seq_form.cgi) and then flanked with target-specific primer binding sites.  The size of each synthetic control was designed to match the actual size of the target. Two to four of these synthetic controls were integrated into a pUCIDT plasmid containing an ampicillin resistance gene, creating a final total of 17 plasmids. *E. coli* DH5-alpha was transformed with these plasmids, and control stocks were stored at -80 °C. Subsequently, plasmids were extracted from overnight *E. coli* cultures in lysogeny broth (LB) containing 100 µg/ml ampicillin using Qiagen QIAprep Spin Miniprep Kit (Qiagen, Cat no. 27104) and plasmid DNA extractions were normalized to 5 ng/µl for further use. These plasmids were pooled and used as a positive control for our assay.
Each table below lists the primer binding sites for each target and each fasta sequence represent a plasmid with synthetic controls. The target-specific primer binding sites flanking the synthetic sequences for each target were highlighted in the same color.

| Organism | Gene | Primer name | Primers (5'→3') | Size |
|---|---|---|---|---|
| ***Coxiella burnetti*** | transposon-like repetitive region | Trans-1-F | TATGTATCCACCGTAGCCAGTC | 687bp |
| | | Trans-2-R | CCCAACAACACCTCCTTATTC | |
| | | RC | GAATAAGGAGGTGTTGTTGGG | |
| | com1 | CoxF2 | ACYGCAGGCGTGGCGATAG | 689bp |
| | | CoxR4 | TGAAGGTTTTGTTGTGAGGTGGC | |
| | | RC | GCCACCTCACAACAAAACCTTCA | |

>Bovreproseq_synthpos_Coxiella_burnetii_1400
TGAGTCTTCGAAGTCACTATGTATCCACCGTAGCCAGTCCGTGATCGGGCCAAACGACGCTATTTCTTGGCCTTACTGCAGCACG
GGGTCCCTGACCATCTTGGAGTGAGTATGACACACAACTGGCGAATTCAAATGACCTTGCTAAAATTACCACTACCGTATCAGAT
TGTTTCGGAACCTTTCGAAGGTAATAAGAACCCTGACCACCGTTAATACTGAAGGTGCAACCATCGACAGATGACTCTGTCAAA
GTAAAAGTCCTTTAAAGGTGCCTCGTTAGTTCGAGCTTCGAAATGCCACAGCTATGGTAGGACAACGCTGTCGTCTCGAACGCG
GTCTAGCAGTCTCCTCACCAGCGCCTTATGCTTGATTCGTTCACCGACCTTTGTGTATCCAGCTTGTAGCCTAGGCTTTTACCCCG
TACGCCTGTACATGGTGTGTCCTAGGCTCCTCATACCTATAAACGACGTTCACAACAATGAAACCGCGACTTAGATCTCCTGTCA
CCGATTATTGCTAACGTGTGGCATGGGAGGTACAAGCCGAGAACGTGTAATTATCGTTCTGACCCGGATCACTTAATGAATAAG
GTCATTAATCATGGCTCACCCTTGATAGCCCGGCCATGACCCGTTGATAAGTCTGACCTCGAAACAGAGGGATTCCTTTAGCATA
AGCTCGAATAAGGAGGTGTTGTTGGGGGCCCACAATTGCACCCAAGTGAATCCACCGCAATCATTCTATCCTGCATCCCGATTT
GGCGGGACACTGCAGGCGTGGCGATAGAGCCGCAAATTTGTTGGCTCCATTTCCAGGCTGCGGGACCACTCTTGAAGTGCTAG
GAGGCTCAGATGCTCTTTATAGGCTGACAGACCGGCGGCTGGCATCCCGTTGACCCCCTCCGTTTCAGTGGTGGAATTAGTGGT
AGTGTGGAGGCTGAGTAAAAGGATTGTATGGAGGCGACCGGCGACTCGCTCAGTCACAGGCCATCACACTATTACCAGTCATG
CACTAAGAGCGTCTTCGGTGCTCGACAGTAACTGCCTGATCGCCCTTCCCCTCTCTTTTATAAGGGGGATGGAAGTCCGTAGTGT
TGATAGCCGCCCGTAACAGACAAGAGCATCGGCGGCCTGCCCGATTCCTTCTACAGTAGGTTTCTTCGTTCGTGATCACCTGCCC
CCGCATCTTGATTTACCTATTTAGAATCTCACCACCGGTCGTCTTAGCCTCGCCCTCATCTGGGGTCTGTAGTGACTATCCAGAGC
GTCCTTTGCCAACCTCTTGATCCCTAGGCTCAGATGGTACGCTTTACATGCCGACTCCCGGAGTTTTGCTCCACGGCTCCTCGCAT
GTGTCTGCGTGTGTCGCCTTAAGCCTTCGGACGCCCGCACGACTCTAGACCGTGCATACTGCATCTAAAAGCGACTGAAGAGAG
CCACCTCACAACAAAACCTTCAATAGCTACGGCATCTGTAGCGCAC

| Organism | Gene | Primername | Primers (5'→3') | Size |
|---|---|---|---|---|
| ***Chlamydia abortus*** | Cabortus Helicasegene | Clone8-Heli-F | TGGTATTCTTGCCGATGAC | 475bp |
| | | Clone8-Heli-R | GATCGTAACTGCTTAATAAACCG | |
| | | RC | CGGTTTATTAAGCAGTTACGATC | |
| ***Listeria monocytogenes*** | Transcriptional regulator | LM1-F | GCTTAATAACCCCTGACCG | 260bp |
| | | LM1-R | AATCCCAATCTTCCTAACCAC | |
| | | RC | GTGGTTAGGAAGATTGGGATT | |
| | hlyA | hlyA-F | GCAGTTGCAAGCGCTTGGAGTGAA | 456bp |
| | | hlyA-R | GCAACGTATCCTCCAGAGTGATCG | |
| | | RC | CGATCACTCTGGAGGATACGTTGC | |

>Bovreproseq_synthpos_Chlamydia_abortus_Listeria_mono
TAATTATAACCGCTCCCGACTGGTATTCTTGCCGATGACTGAGTTGGGAAACGCATTAGCGTCGAATGCGTCTTGCGGGAGCTC
GCCCCTGTTCAATCTTGGTTGATGTCTTTCTCTAGGTCCTAGTGTCAGTTCGAGGATCCCCATGGGTAGATTTATATACAAGGCC
GTGTTAACGGAGCAAAAATAGACACGTGAACCCACCCGGTGACTATGAGGTGTCGGATACATGTTGACCTCCGTTGCTCGCATT
TGTGCTATCTTCGGGGCTGACCTAACTCGGGTGTACATCGGTGACAATCAGCCTTAGGTCAATCAGACTGTCTAGCTGGTGCGC
CGCAGAACTCCATAATAGCCAATAGTATTAATGTCTCTAGAGAATTTCAGCCACGCGCCATCACGTAACTATCCCCACTATGCGA
GGATACTCATAATGAGAGGTTAGCGCCCCGGGGTGAGTCCTCTTCTTACCTGGATCGGTTTATTAAGCAGTTACGATCCGGACG
TACGAGGGCGCGGCACAGGATGGCGCGCTTAATAACCCCTGACCGGTAGTGCCGCTACTGCATTCACAAGTGCTCAATTGCATC
CTCCCAGATACTTTTATTACCAGGTCTCCCGACAAGTTACTTCGAGTATTACTGTTCGCATATTCTAGTTGGTCTGGCGCGCGGA
GTGTGCGGCCAGGCGCCGTCGAGCGCTCGTTAAATGTGTCTAGCGTGGAAAAACAGCACTTGACGAAGTGCCATTATCTCTCG
GGCAGCTGAGAATGTGGTTAGGAAGATTGGGATTCCAGCAAAGGAGCCCCAATTCTGTGGATGTAGCCCTCCTGGTGAATTCC
GTGCAGTTGCAAGCGCTTGGAGTGAACCGAGGGTATGGAGTGGGTACGGGTAAACTCTTGCGACCGTGAGTCCGGGTGCTTA
AAGACGGGGAGATCGTGCAGACCAGTTTATTTAATATCTTACTTTACGCGAGGCTACGCGGCCCGGTCTGGCGGTTGCCATGCC
CATCACAGTTTGTCCAAGCTTGGATGGCCTTGACAAGTGGTATTCGCTATTAGCATGGGAAGCCCCATAGCTTCCTGCAGTGAT
GACAAATATAGTCCATACCTAGCTGAGCATGATGCGACCTCCTCAACGTTCTCCCGTATCTATGAGGATCACGGTGATGTTGTTA
CCACAGGCCTTCAAGATTTATGCTGGCTCAAGGCTCCCTGCGACCAGCAAGTCGCGTCCGTGAGAGCGATACGACCCCGATCCC
GTGAACTGGACCTCCGATCACTCTGGAGGATACGTTGCTCCCTCGCATAGGAATGACGTTAGT

| Organism | Gene | Primer name | Primers (5'→3') | Size |
|---|---|---|---|---|
| ***Ureaplasma diversum*** | 16srRNA | UDF1- | AAATGTCGGCTCG**C**TTATGAG | 311bp |
| | | UDF2- | AAATGTCGGCTCG**A**TTATGAG | |
| | | UDF | AAATGTCGGCTCGMTTATGAG | |
| | | UDR | TATCGATAGATAAATTAACTAGCG | |
| | | RC | CGCTAGTTAATTTATCTATCGATA | |
| ***Ureaplasma* spp.** | 16srRNA | UGPF- | GGATGAGGGTGCGACGTATC | 642bp |
| | | UGPR- | GCGTTAGCTACAACACCGAC | |
| | | RC | GTCGGTGTTGTAGCTAACGC | |

>Bovreproseq_synthpos_Ureaplasma_diversum_1400
ATTTGGTTTACGCTTGTGAAATCGGCGGAAATGTCGGCTCGCTTATGAGTCTGCCCAACGCGATCCGTACGAGGTCAGATGACA
GGCCCCGTCTTCGTTACCCGCCTATCCTGTCCACCATATGATGCTTCCGGTAGAGCGGGTGGCGTTAACCACAATTAAACGATTA
GGTCGCTTGAGTACGTACCCCTACCCGCGGCTTGAATACCTAAAAAATGCCCCTACAGACCTGTAACGCGATGTAATTTCGCAAC
TTATACACGCCCATCAAAAACAATAACCAAGAGTCGGGCTGGTCGAGTGACCAAGTGCCGACGCTAGTTAATTTATCTATCGAT
AGTGGGCGGGCCGCATGGTGTGCACACATCCAAGTCTTTTCCGCAGTCCCTAGCGCCGTCGAGTATCGCGGTATGACCTGGAT
GAGGGTGCGACGTATCGTGGGAAGGGTGCGTGAACAGAATTTAAAAGGGACCCGACTTATATGTCCCAAAGTGTCGCGCGGA
TGGATCTTAGTCCAGTCGTTCGGACCGCGAGGATGGGATAATCATTCACGCCCTGCCCACTTGTTCGCTCCGTCCTGCTTATTTA
CCCTTACATCTACCCAAAATACAGTGCGGGGAGTAACACTATCTGAAGAATTTGGAATTTTGGGAGTAGGCAACCGGCACACGC
CCTAACCGTAGTGTATATTTACGGATTCGGGTCGATCGGCTACGATATTTTAGGACGCTTACCCGTGTCCCACTGTAGCAACTGC
AGTGAGTTGGACTTGCTTAGTAGGAGGTCTTAGCTCAATCACACCTGTTTGACCAGCAATAGTTAACATCTCGACACATGGCTCA
ACATTATAATTTGATTGCTTTTAAAACATACGACAGAAAGTAGTTCCGACTTGATGATGCATGACCTTCTGTGAGTACTCCAGCTA
CGAGATGCCTTTCGCCCAGAGTGCGAGAGTTTTTCGTCGTACAAGGACTTACTACGCCACGTCCAATCGAGCAACATATGCGTC
CAGTGTCCACCCTATGAACCCGAGTCAGTCGGTGTTGTAGCTAACGCAGTGCACCGACCTTATGTAAGTAAATAC

| Organism | Gene | Primer name | Primers (5'→3') | Size |
|---|---|---|---|---|
| ***Leptospira* spp.** | 16SribosomalRNA (rrs2) | rrs2-F | CATGCAAGTCAAGCGGAGTA | 541bp |
| | | rrs2-R | AGTTGAGCCCGCAGTTTTC | |
| | | RC | GAAAACTGCGGGCTCAACT | |
| | LipL32 | Lau01 | ACTCTTTGCAAGCATTACCGC | 660bp |
| | | Lau02 | AGCAGACCAACAGATGCAACG | |
| | | RC | CGTTGCATCTGTTGGTCTGCT | |

>Bovreproseq_synthpos_Leptospiraspp. _1400
CCCCAATCAGAGTTGATATGAGGCATGCAAGTCAAGCGGAGTAGGTTGACCACTAGTAGTTTCTAAGTGGATACCCACCGGCA
GCATTGTAGGGTACACGGGGTACGGTAGGTCTACGTACGTAACCCCGGTGTTATTAACTGGACTGTTCAATGGGGACGATCTAT
TTTCCAGCGTCTCATAACCGGACTAACTCAGCGGGGGCACAAGGCACAGAGTTATACGCGTATCTAAGCTGTCGCGTCGCAAGG
CAACTTAGCAGATTACCGGCGTAGTGGATACGTTAGAGGTAACGGGCCTTAAGACGGCGCGTATGGTGATACACGGCGACAGA
AGTCTTTGGCCAGGCGGCCATCCTTCTAACTAGGTAAATTCGACTGAGTCTGCCAAAAGGAGTTGCGTGTCCTTCCTGACCGAG
GGGTTGTTCAACCATGCCCGTACGCGGTCTACCAGCGTGTCCTTCAGCGACTGCAAGCAGTCAAAGCTTCTCTGCGCGCCATTA
ATTGATTAAAGAAACGCCAAGTACATGAACACGGTTCCAACAATAGAAAACTGCGGGCTCAACTTATGATGAAACGGAGGGAA
ACTGAACCTAGCAGCGCCCAAGACTCGTGACTCTTTGCAAGCATTACCGCTTACTCGCTTGTGGTTGTGTTGCTAGTAGGGTAG
GTCAATGCGTATGCGGAAGTCGCTGGGTATGCTCAATCAGTTCAAGGCCTGTGTCTTGACGTTTTCACGTCACACTGAGGTGAG
CGGCTGATAGAGCGAATGCTCAAATGTCCAAGAACGACCCTGCGCGGTCAAAATAATCCTATAGAGCTTAGGATGATACGGCA
CATTACCGGGGGGAAGAGCGTAGGACGGCGACCTTGCTTTTATCGCGTACCAGATATCTTAGGGTCTCGACAACTAACTATCTG
CGCTGTATTCGTAGGGCATGTCTGCAAGTATACACAATGTGAGCCGCAGAAAGCCAGCACGAGGCCCGCTTAGTTAGCGGCTG
TGGGTACTTGGCTACATAGAGGTATAGGCTGTACAATCTGGTATCGATGGTACGATGTAAACCAAATTTCGGCGGTGTCAGCGC
TCCGGCCGCGGACCTCTGCAACTCAGAAAGCATCAGATGTATTACTCTTCAATGAATCGCTATGGGGCATCTAATGACAAGAG
GTATAGAAAACATAAGCACGGGCTCACCGTGCGCAATGGCGATGGTATATCCCCTACTTCGGAAGCGACGCCAAGGCAAGCGT
CGTTGCATCTGTTGGTCTGCTTACTCCATCCCGGGGGTATTACTGGTGGCCTAATTT

| Organism | Gene | Primer name | Primers (5'→3') | Size |
|---|---|---|---|---|
| *Neospora caninum* | pNC-5gene | Np21 | GTGCGTCCAATCCTGTAAC | 328bp |
| | | Np6 | CAGTCAACCTACGTCTTCT | |
| | | RC | AGAAGACGTAGGTTGACTG | |
| *Tritrichomonas fetus* | ssrRNA | TFR3 | CGGGTCTTCCTATATGAGACAGAACC | 347bp |
| | | TFR4 | CCTGCCGTTGGATCAGTTTCGTTAA | |
| | | RC | TTAACGAAACTGATCCAACGGCAGG | |

>Bovreproseq_synthpos_Neosporacaninum_Trich_fetus_1400
AGCTCAGAAGCCTTAAGTCTATGTGCGTCCAATCCTGTAACGGAGCCCGTATAGATATATCCACGCTACGTTGGTCCTAGCAAG
GGGTCCCTAGAAGTCGTGGTTTGAAGCTTAGGCTAGTCCCTCGCGTAAAAAGAAACCTACTCGTGCCGGCTGAGTACACGAGC
CATTTCTGCCCAAATGCTCATGTCGGCAAATATACCTCGCCATGCTCCTCTCGTAGTGCATGTCACCTAATATGAATATTCCGTTA
CCGTCCTGCAGTGTTTTTCTCCGGTTAGGAGACCAAGTGAGCCAGCGTAGCCTACATTGCCTAGCACTCCAACGCTCGAGAAGA
CGTAGGTTGACTGTGTCCAATGGGTACTCGCAACTGGATAAGTGGAATGGAGCCTGATAATCGGTTCTGAGGTCGCGAATTTTC
ATCCGGGTCTTCCTATATGAGACAGAACCATATGACTCCCCGCACCCACATGACCCTGTATACTGCCTCGGCTGAGGTGGGTTTC
CAGCTCTCTTGGCAATGGACGCTAACGAGTACCGAGATTGCAGTCGTTCAACTCAGGCGCTACGGGCCCGCTTACCGTAATCGG
CGGGTCGACTGATGTCCCTGGCGGCCGAATAATCCCGTTAGCACAGTTTCGGCTCTAGGCTTGGCGCAACCCTTGCGGTCTCGG
TGAATTCCCGTGTATTGTGCTCGATCTCCCCTCCAATCCACAAGATACCACATGATCTATGATTAGGTCACTTTAACGAAACTGAT
CCAACGGCAGGGTCAAGAATGCTTCAGTGT

| Organism | Gene | Primer name | Primers (5'→3') | Size |
|---|---|---|---|---|
| **Bovine Herpes Virus Type1 (BoHV-1)-IBR virus** | gBN-1 | gBN-1- | TCTCGACCGGGGACATTATC | 385bp |
| | | gBN-2 | GCCTCTTCGATCACGCAGTC | |
| | | RC | GACTGCGTGATCGAAGAGGC | |
| | gE-1c | gE-1 | GCTTCGGTCGACACGGTCTT | 268bp |
| | | gE-2 | CTTTGTCGCCCGTTGAGTCG | |
| | | RC | CGACTCAACGGGCGACAAAG | |

>Bovreproseq_synthpos_IBR-BoHV-1_1400
AAGCTATAGATCCCGGGTGTCTCGGGCCGATATCTCGACCGGGGACATTATCTGATCTAAGCTACCCACTCACCTGCAGGCAAT
ACATCCCGGGTGATACGAAAACTTCACCCGCGGCAAACAGGACTGCCGCTAGTGTTTAGTGGGGAGAGAGCAACGAAACACAC
GGTTTTACGTTCATTGTTCCGGCAGTACGCATCATTGTAACGTGCGGCTCTCATCAGAGTTGCAGTGCGCAAACCCGCGCTCGAC
ACCCTACCGCTACAACCGCACTTCACGAAATCCAAAAACCTACTTGGCCGCAAAATTATGTATCGTAGATTGCGGCGGTTACGGC
TAGTAACGTAACCAATATCAGTGCCTCGGTCTTGTCCTGACAAGAGCCAAATTAGCTAGTGACTGCGTGATCGAAGAGGCTTAA
TGGGGTAACAATGGAGCACGCGCTGTAACTGAGGCTTCGGTCGACACGGTCTTTTTATTGTAGCGCACGATACTGTTACCCTAT
TACGTAGAATGGTACGCAAACTGTGGAACGTACAGATGTCGAGGCTCTGCTCATCGTAGTAAGGTGGTGCATTGTTGTCACCGA
GAATGGTACAGCACGATACGTTGATCGTTGTCAGGGCGGTGGAGCGCTTGAAGGAGTGCGTAAAAATTCAACAAAGGCCTACT
CACGCTAGTGAAACAGAGGCTGCCACCATTCGACTCAACGGGCGACAAAGCTCGGGACCATCCAACCTCGTCTATCTGC

| Organism | Gene | Primer name | Primers (5'→3') | Size |
|---|---|---|---|---|
| *BVDv* | 5'UTR | PF1 | ATGCCCTTAGTAGGACTAGC | 285bp |
| | | PR1 | ACTCCATGTGCCATGTACAG | |
| | | RC | CTGTACATGGCACATGGAGT | |
| | Npro | B32(Forward) | CCATCTATRCAYACATARATGTGGT | 441bp |
| | | B31(Reverse) | TGCTACTAAAAATCTCTGCTGT | |
| | | RC | ACAGCAGAGATTTTTAGTAGCA | |

>Bovreproseq_synthpos_BVDv_1400
CTATTAGTACCAGCTTCACACGATTCAGTTAGCCGTCGATCAGATGCCCTTAGTAGGACTAGCCTACCTTTACAATCAATACTATC
GTCCTATACTCTTCACAAATATGCGTTGTACCAGGGTGGACGTTTCTTCAGTGACGCGTATTCCTACTAGGTCATATGGTCTCCAT
TCCGTGCGCTAGATACGGTCGAGATGGCGTGATCTAAAGTTGTGCTAAGGACAGTATGGCTTGGCGCGTCGGAACAATGTAGT
TTTAAGAGTACTCATGCGGAAGTAGCTTAAGCTTCAGTGCAAGACTTTACTTCCTGTACATGGCACATGGAGTGACCTAAGGATT
ACTGTTGTGGATGACCCTGCGTGAATAAACG**CCATCTATACATACATAGATGTGGT**GCGCAGAATCACCGCATCATAACCGAAG
ATGAAACAGCACTTGGGGGCTCCAGAAACGTAGGCCTCCGCCGGACTGCCGCGATTTCACAGTGCCGATTACAACATATCTCTG
ATCCTTAGGATGTGTGCCTAGGACCCTTACTCACTTGAGCACAAACTGAACATCTGCGTTGCTAAACGAACGGACCACGTTGGC
GTTACCTTTCCGCGACGACGCAGCCGCGGTGACTGTTTTAGCCATCTCGAGGTCGCGATAACTTGGTTGTGGGGAGCTACTTTG
GTTCATACCACGCAACACCGTACAGCGCGACCAGATTCCACAGCGCGGGGATTAGGACGGTAAAAAGGTAATCTGGAAGTTTT
GTAGGCAGGTTGATCCTTGTAGAGGGGATAG**ACAGCAGAGATTTTTAGTAGCA**GGCGAATTATGATACCTTCACCTAGATCAGC
ACAGGCACGC

| Organism | Gene | Primer name | Primers (5'→3') | Size |
|---|---|---|---|---|
| *Campylobacter fetus* | nahE | nahE-CFETSpp-L | GGTTATTTTTTATAACTGTAGGAATGCAGAT | 390bp |
| | | nahE-CFETSpp-R | GATCGCTTAAATCTTGTACTTTTAGCTTTT | |
| | | RC | AAAAGCTAAAAGTACAAGATTTAAGCGATC | |
| *Campylobacter fetus.* Subsp. *fetus* | sapB2 | CF-F | GCAAATATAAATGTAAGCGGAGAG | 433bp |
| | | CF-R | TGCAGCGGCCCCACCTAT | |
| | | RC | ATAGGTGGGGCCGCTGCA | |
| *Campylobacter fetus.* Subsp. *venerealis* | *virB11* | *nC1165g4F* | AGGACACAAATGGTAACTGG | 233bp |
| | | *nC1165g4R* | GATTGTATAGCGGACTTTGC | |
| | | *RC* | GCAAAGTCCGCTATACAATC | |
| | *parA* | *VenSF* | CTTAGCAGTTTGCGATATTGCCATT | 142bp |
| | | *VenSR* | GCTTTTGAGATAACAATAAGAGCTT | |
| | | *RC* | AAGCTCTTATTGTTATCTCAAAAGC | |

>Bovreproseq_synthpos_Campfetus_CFF_CFV_1400
AAGACAAAAGGGATTAGTGGGGTTATTTTTTATAACTGTAGGAATGCAGATTCAGACACAGGGTTGGCTTATTTGCGTCTGGTA
TGTCTTTAACAGCGATGCTATTTAAGGCACGAAGGGAAAGCAGCAAACCCTTTCTAACGCTAAATACCTGTGATATAGCCAGCTT
ATACTAGTGAATGTCCCCCCTTGCTAACGTCGTATGCGTTTTGCGTCCATTGCGATACCTACAAGGTAGCGCTTTATGCTTCAAAT
TCTGGGGAGCGTGGCATACGTACGATACTGAGACTTTATCCAACAAGATGGGGGCTCCACGTGTTTGGTGATTAGAAAGGAGC
GACTGGATGGCAAGAGTTACACGATATAAGGCCGGTACCATCTAAAAGCTAAAAGTACAAGATTTAAGCGATCAGATGCCACAC
CATGAAAAGATGCGTACGTGGAGGACATGTGGCAATTGTTCGCACGGTTGTTATGTTCAGCAGGCTGAACAAGACATGGAGGG
GTATTTTCAAGGCGCATTAAGTTAGGTTCCGTCGTTACCTGCAAATATAAATGTAAGCGGAGAGGCGGCTCGTTCGGTCTACAG
ACTATGAACCGGCAGGATCTCCTAATCTCCCTCAGCTCCATCGACTCCGAATTTCGTAAAGTTCAGACATCAAGCGATTGCAGGG
ACCAAGATCATCATAAAGCAATTTTCAGTAGACCGTCGTATTCTAAACTAGAAAATATCCTCCCAATGGATGGATCCCGGAGAAG
ACAGGCGTAACCGCATGGAGGAAGAGTCCTACATCCTCACTGTTTCACCTCATTTACGACCCAACGTAAGGCAGTAGGCAGAAG
TTACCCTCCCCTGTGCAATCAAAGCAACGCTTATGGCATAAAGAAGCCGTTACCATAACGATACTCGTCTTAACCTTTGGCCGTTT
CCGGCCCTGCGGCTTGAGACCACTTACCGGATAGGTGGGGCCGCTGCATATCATAAATGTATTGACAGGATCGGTGCGATTAG
AACCTGGGTTCCCTTTTAGCCGCACTACACGGACCAACTCAGTGAAAAGAGAAAGGACACAAATGGTAACTGGTGACGTGGGC
ATTCCGTCACCTCAAAATGGTATGCAGATGCGGTGGGACTAAGAGCCATTATCTATGATGTGTGCTGCGAAAGCAGTCTACGCA
ACCGAGGCCGAAGGGACTGGCAATGCAGGTCTGCCAGGTGCGCGCGTGTGGTACTAAGGTGTCTTACTTTATTAATAGTCGCG
TACGCAACAGCGCGCTCTTGGAGCAAAGTCCGCTATACAATCTATCACCGTGTGATCTCGGGAGTGACGAGGGGGCATTTTTTA
TTATGAAAGAGGCAGAATGCTATCGCCGGTTCAGGGGCATGATGAACGATGCCACAACATGCGCCTTAGCAGTTTGCGATATTG
CCATTCCAACCTGTACGTGGTTGAGGATAGAAAGATAACTCAATGCCCGGCAATCATTCGGGAAATGACCGCAGCCTCGTATCG
CTGGATGGAACAAAAGCTCTTATTGTTATCTCAAAAGCCTCGCGCACCGTCAACTACAACCCACCACT

| Organism | Gene | Primer name | Primers (5'→3') | Size |
|---|---|---|---|---|
| ***Campylobacter fetus* MLST** | *aspA* | *aspA-F* | CCTATGACTTTAGGTCAAGAG | 477bp |
| | | *aspA-R* | TGTAGCTAGAGTACGGCAAG | |
| | | *RC* | CTTGCCGTACTCTAGCTACA | |
| | *glnA* | *glnA-F* | GATGGTAGTTCTATAGACGC | 477bp |
| | | *glnA-R* | CTTCCGTTATCTCCATAAAGC | |
| | | *RC* | GCTTTATGGAGATAACGGAAG | |
| | *gltA* | *gltA-F* | CGATATAGCGTGGCTAGCTG | 402bp |
| | | *gltA-R* | AGCGTGAGTAGATCCTACG | |
| | | *RC* | CGTAGGATCTACTCACGCT | |

>Bovreproseq_synthpos_CampfetusMLST_aspA_glnA_gltA_1400
CCACCCAGGCGCCTATGACTTTAGGTCAAGAGGAATGCATGCCCATTGCACACGGGCCCAGATAGTACGGAAGGATTTGAGGA
GCTCAATTCTGATTGAACTGCGCCGTTAGCGCAGATCGTACTCCGTGTCTATCTCCACTACCACATGTTCCGAACTAGACTGACT
GATTTACCAAGAGGAAGCAATATGTTATAAACTCATTGTGTTGAATAGCGGTAGGCGTAGAACACAGTCAATTGATTTATTTTTT
ATTGTGCCTGTAAGGCTTGCGTAGACTACAAGTCGGGTCTTTCCAGAACGGTGCGTATCAAACTTCAGCGCAGACTAGGCTACA
GGAAGAAGCGAAAGACCCCCCTAACGCAAACTGTATGCAGTCTTTGTATGCTCACACTCCCTCGAAGTCTCACAATGTCCGGGC
GGTTCCCGAGTTAGAATCAAATCAGCGGTCATGTACCGTGGGCTTCCCTTGCCGTACTCTAGCTACACTCGTCTGAGCTAATTCT
GAGTTAATCGATTTGCACGTTACATCAATAATGTGATGGTAGTTCTATAGACGCGATTCAGGTCACCACTCCATATGTTACCCACT
CTCGTGAAGCGTCAAGCAAGAGTTGTCAATTCGGTATAGGCTCGTGTGTCCGTGTTATTGCCGCATGTAAAGCGGCAGCGCCGC
GGTTGATTGAATAGTGTGCCGGGGCTGGCAGAGTGGTTGGAGCGCTCTAGTCATAAGGATTGACATGCGATAATAGGTCCAAT
GCAAGGTGTTCGGACGTTATCTCATTCCAATGCAATGCGTATCACATGCTCGTGGTGAGTGGGGCGATGTACGACCTGTTGGTG
GCATTCATCCCGGGGTTAAGACGCTCCCGTGTCCAACCCCATCCCTTAATTGAGTTGCGGTTGCTTTCGCGCCCAAAAAAGCCGT
CCTCCAACTCCGGGCAGATGGAGTAGGTTTAACCAGATTCTCCTCAGAAGTACCGTTACGAGCTCGCTGCTTTATGGAGATAAC
GGAAGCAAGCGTACCCGCCCGTCGGCGAACCGTGAAGCGATATAGCGTGGCTAGCTGGCTTACAATACTACGGCAAACTGGCA
AGGTCCCAACAACTCTTAAGCCCAGCCGTGCCGGTAACGAGACTCGACGGCGTTCAATCATCGAGACTGGCTCTCTCTTGTTGG
TGAGTCGACACACATGCGGGATCACTCGTCAACTTTCACGCTACTTAGCCTCTGGGACCTTCCCAGGTGTCCAATCCACCTTGGA
TGAGGGATATGCTCGACCTCGGGCCTCGGAGATTCTTGGTGAAGTGACGGTTCTGACAGCCGATGTTATGTGAGTAGGATTTTG
GTCTTTGCCGGTGTTGCATATCCTGGACGCGCTGAACTTAGAGGCCCCCGCGCACACGTCAATAGAACCCGGGCCCGGCCGTCC
CGTAGGATCTACTCACGCTCAAATTATGCACTATCTAGATCTACGCGGCTGAAGAAGTAATTATTGACTAAGGTGATGAATTACT

| Organism | Gene | Primer name | Primers (5'→3') | Size |
|---|---|---|---|---|
| *Campylobacter fetus* MLST | *glyA* | *glyA-F* | GATAAAATACTTGGTATGGATC | 507bp |
| | | *glyA-R* | CCCTCTGTTTATTAAGACTTC | |
| | | *RC* | GAAGTCTTAATAAACAGAGGG | |
| | *pgm* | *pgm-F* | AGAGTTGTTTTGGACGTTGC | 501bp |
| | | *pgm-R* | GTAGCTCATCAAGAGGTCTC | |
| | | *RC* | GAGACCTCTTGATGAGCTAC | |

>Bovreproseq_synthpos_CampfetusMLST_glyA_pgmA_1400
GAAGCTTATGTTCATGTGATAAAATACTTGGTATGGATCAACACATCCCCGTTCGAGGTCAAAGGGCGTGGTTTTCGGAGTCTG
TGACATCTTTCGCCGGTAGTCATTCGTCAGCCATATGGCTACTCTGTCCTCGCCGAACTGTGTATCCCACCCCACGAGACGCAGA
TCATGATCCCTCTTGGGAGGGAGGTCACATGTCTGAACCCTGTACCGGATCGAAAGACTAAAACATAGAGAAACTTTCGCTGGT
TCCTGCGCACCATCGCGACTTCAAACGTAAGCTGTTAAATCCGTCGAAAAAATAGCTCTTGCGAGGCCAGGCTGCGCGGCCTGC
ATTGGGCATATTGTGAATCCGATAGCTGTGAGCTTGACATTTTTCCGAAACGTAAGCACTCGCTCGGCGTTCGCCGGGACGGGT
AAAGAAGTGGGTCGATGACCAAGACTGCAAGCAGGAGGACAGGCGCACATCTTTCATTGGGGTTTATTGGCGGTAGTGCTTCG
AGATCCACTCTTCTCCGGAAGTCTTAATAAACAGAGGGTTGGGTGTTTTAACTCAGCCATTGCTTCTACGGCTACTTCGGGCGTG
CGCGAGGCTACTTCGACTGGAATGTAGGAGAGTTGTTTTGGACGTTGCTGTCCAGTTTTGGAGATTCTCCGCTGGCCGGGGTG
GCACTACACTGTTCAGTCCTGTGCTCACCAGATGTGCATCATCGGACTAGGTAGATCTTGGTCCCGGGGAACCAGCGTAGGATT
CTGGTTGGTGCTGCTGACCGAGGCGGCCATACCATTGGGTCTAAAAAGACACAAGGTGCTCGTATTTTGCGTCAGGTGATACTC
GTCGCTCCCACTTGGCGTTGTACACACGCAGCATATCGATTTGTCAGAGGTGTCTCGAGGTCATATACCTATATATTTAGGGGTT
GTACTGCCACTGCTAGATTACCGGGCCTTCTCAGAGCCCGAGTGAGTCATGTGCCAAAGTTCTGTGTTTTTCGCGGGAGATATTC
GTGGTCCCACGGTTCTCGCTAACGGAGGACTGACCGCCGCTTAGCGTCCACCGTAAAAAGATGCATATGAGCGGGTAATGGGT
GGGCAGTTTCTGAGACCTCTTGATGAGCTACTTTGGATGGGGATTATAGGTATCCAA

| Organism | Gene | Primer name | Primers (5'→3') | Size |
|---|---|---|---|---|
| ***Campylobacter fetus*** **MLST** | *tkt* | *tkt-F* | GAGATAGATTGGTATTTAGCGG | 459bp |
| | | *tkt-R* | GTGACTACCTTCTAAATCTCC | |
| | | *RC* | GGAGATTTGGAAGGTAGTCAC | |
| | *uncA* | *uncA-F* | AAGAGTACGGTGCTATGGAC | 489bp |
| | | *uncA-R* | CTCTCATCAAGATCGCTTGC | |
| | | *RC* | GCAAGCGATCTTGATGAGAG | |

>Bovreproseq_synthpos_CampfetusMLST_uncA_tktA_1400
CGATACACAATGGATCTCTGACTTCCGGAGGCAGAGATAGATTGGTATTTAGCGGTCTAATGTCCAAAATAACGCGGATCGGCA
CATTAGTAGTATTTCTCAAGTTGGCTCAACGACAGAAATTCGCCGGCGAGGATGTGGCCTGTATCCAGCAATCGGCTCGAGAAT
TTTGACCGCACGTCCTAACCCCGTCTCTCAGTATCGACACCTCGCTTACCAAACCATCAGGGAAACCCTTAGACAAAGGTAAAGG
ATCTCCCCGTTGCGTCATCAGTCCCAACAGATATGATGTGCGGTTGTCCGGACTCTATGCGGGAGAAACTGCGGTGTGCTTCCG
ACCCTTTGCTGCTGCGTGCTTAGGGGGACTATTAGGTAAAAGTAGTCCAGATTTAGGTACATATACATGATCAAGTTCAACCGAT
GCCCTACAGCCATAGCAGTGCCTAGTGCTCAGGCTCCATGTTACTAAGCCGTGGAGATTTGGAAGGTAGTCACTGAACAAATGG
CGTGCTGCCCTGAGTACTGCGACGGTCTCGCGGAGATAAGAGTACGGTGCTATGGACGTGCTATTCGGTGATTAGAAGGGTCG
GGCCTATAACGAGTGTACCAATGTATACAGCCGACATAGTTGTCAATGAGCCGAGCGTCATCTGTAATATGCGTTACGGCCCTG
CGGGCCACGTCCTAGAAGTATGTGGTTCGCTAGTGCAAGGTCGTGATTTAAACGATCGCCCACTTCCCGCAGCACGCCTAGTAG
CTTTTATTCAATGCTAGGGGAGGCACGTCATGCGGAAAGCCGTATGTTTCGTCGTGTTACTTCTACAGTTTACTTATAAGTGCGG
AGCCCTGTTCGTAGTGCGGACAGCTTACAATGCGGTTGGAAATGCGCTGGGACAAGAAGTACCTACATCGAGGATGGCTACCG
ACTATGAGATAGTCCAATGTTGCGAACTTTGGGCCATGTCAAGGCGCAGGCGTTACCTAGGAGATCCGCGCGCGGCAGTATCC
CAGCGCAAGCGATCTTGATGAGAGACTCTCAGTGATCGAGGATGCCTTAC

| Organism | Gene | Primer name | Primers (5'→3') | Size |
|---|---|---|---|---|
| **Internal Control Enhanced green fluorescent Protein** | IC-EGFP | >Egfp-F | GACGTAAACGGCCACAAGTT | 550bp |
| | | >Egfp-R | GGGTGCTCAGGTAGTGGTTG | |
| | | RC | CAACCACTACCTGAGCACCC | |

>Internal control enhanced green fluorescent protein gene
GACGTAAACGGCCACAAGTTCAGCGTGTCCGGCGAGGGCGAGGGCGATGCCACCTACGGCAAGCTGACCCTGAAGTTCAT
CTGCACCACCGGCAAGCTGCCCGTGCCCTGGCCCACCCTCGTGACCACCCTGACCTACGGCGTGCAGTGCTTCAGCCGCT
ACCCCGACCACATGAAGCAGCACGACTTCTTCAAGTCCGCCATGCCCGAAGGCTACGTCCAGGAGCGCACCATCTTCTTC
AAGGACGACGGCAACTACAAGACCCGCGCCGAGGTGAAGTTCGAGGGCGACACCCTGGTGAACCGCATCGAGCTGAAGGG
CATCGACTTCAAGGAGGACGGCAACATCCTGGGGCACAAGCTGGAGTACAACTACAACAGCCACAACGTCTATATCATGG
CCGACAAGCAGAAGAACGGCATCAAGGTGAACTTCAAGATCCGCCACAACATCGAGGACGGCAGCGTGCAGCTCGCCGAC
CACTACCAGCAGAACACCCCCATCGGCGACGGCCCCGTGCTGCTGCCCGACAACCACTACCTGAGCACCC

| Organism | Gene | Primer name | Primers (5'→3') | Size |
|---|---|---|---|---|
| ***Toxoplasma gondii*** | Major Surface antigen | Toxoplasma_gondii_DS38 | CGACAGCCGCGGTCATTCTC | 550bp |
| | | Toxoplasma_gondii_DS39 | GCAACCAGTCAGCGTCGTCC | |
| | | RC | GGACGACGCTGACTGGTTGC | |
| *Sarcocystis* spp. | 18SrRNA | Sar-F1 | GCACTTGATGAATTCTGGCA | 609bp |
| | | Sar-F2 | CACCACCCATAGAATCAAG | |
| | | RC | CTTGATTCTATGGGTGGTG | |
| *Pan coccidia* spp. | 18SrRNA | Coccidia_18s-F | GTTGTTGCAGTTAAAAAGCTCGT | |
| | | Coccidia_18s-R | ATCTAAGAATTTCACCTCTGACAGT | |
| | | RC | ACTGTCAGAGGTGAAATTCTTAGAT | |

>Bovreproseq_synthpos_Toxo_gondi_Sarcocystis_sp_Coccidia
TTGGTTATGGGTTGTTGCAGTTAAAAAGCTGGTGGCCAAATTACTTCTCGTCTTTCCACCCACGTGGATCTAATCGGATGTGGTC
GCTTCTACCCCGTTGCACCCAAATTCGCAATATTACTATATTGACAGGAGTATCGACATAAGGTGTCGGTGTCCGGGAAAAGGG
CGATAGATTGGCAATCTTCGGGCTCGGATACGTTAGCCCTTTAATTCAGACCTATTGAGTCCATGGGTCATCTCCCGCTTGCAGC
CAATAAAGCGGGAGGAAAGTTTTAGCTACCCCAAGTCGTATCATAGCCTGCACTGTCAGAGGTGAAATTCTTAGATCTGAGAAA
GACACTAGTACGATGTGCACTTGATGAATTCTGGCAGCCCTCTCGGCCGGGGACGTGTGCACCGAGATCTTATCTTATTTTAGCT
TACAGACTATCGCACCACTCTCTTGGCTGTAGTAGGTTACACGTCGCACTCGAGGTCGCTCAGGTTTCTTCGGTCGCTGACGTGT
TTATCAAGACATATACCTGATTGGATCGGACGAGACGTTCGGTAGCTAAACGGTTCTGCGAGTAGACAGCGAGAGGCCCCACCT
GATGCAAACCTGATGGATGGGACGTTGGACTCAGTGCCGATCGATTGGCTTCTGTTGGTCTGTACCACGACATCGTTGGGACAT
AATTGCAATAGCGAACTGAAGATGCCGCTCCCCTTTCCCCGCGGCTAGGGGCCAAAACATCTGATATCCTAAACACAGTATAGTT
TCTGCAGTTGAAAGTCTGCGACCGACTTCGTTAGTGATGGCCGCCTAGCGCGGGGGCACGCGGGCTAGAGCGTACCCCCCGGC
TATGCCCGTTCCCACAGTCCTTCTCGGCATTATGTCGGTGCTTAGAGTGGGTACCGATATCACCGGTTGCTATCTGCCTTGGCTA
CGCGCATATTCTAGGCTTGATTCTATGGGTGGTGATATTCTAGGAGGTGGGAAGGTCGACAGCCGCGGTCATTCTCAACTGTTG
TTAAGCAAGCCATGTCCAAGAGCGTACCCTCGTGGAATAGGTGGTAACAATTGTTGCAGACCAGTCTTCTGCTCCGGCCTGCCC
TATCCGTGTATAGCATGGCTCATCAAGGTCAGTCCCATTACATAAAAGTGATACTTTTGTGGCCTGTTTCTGGTAGCCTCTGCGG
CCAGTCCTTTACAAAGATTATAGGTCGCAGTTTATGCAGCTTAGGACATGAGCGGTGTGTAGATCAACAAGCTCCTCCCTCGCCT
GGAGAATTGAGACATGGATTCCTTAGACGATTAAGGTCCATGCTTTATCAACGCTCGGCTGATTCAAGGGTCGTCGCGAATAGT
CAGAACCGATCGCGATTCACAATATAATTAGTGGATATATTGAGGATGGCTCCGATCATCTCGACGAATATTAAGACGAGGCCG
TCCTCGAAGGTAACGGAAAATTGCCTACCCAAGGACGTGGCAGTCACTATCCAAAGCCTGTTTGCTTGCGATGGTCGCAAGGAC
GACGCTGACTGGTTGCCTTCCGTATATGTGAA

| Organism | Gene | Primer name | Primers (5'→3') | Size |
|---|---|---|---|---|
| *Bacillus licheniformis* | adk-adenylatekinase | Blich-adk-F | GGTAAAGGGACACAGGCTGA | 613bp |
| | | Blich-adk-R | TCGAGTAAAGGCTGGGTTTG | |
| | | RC | CAAACCCAGCCTTTACTCGA | |
| | gyrB-gyraseB | Blich-gyrB-F | AKACGGAAGTGACGGGAAC | 518bp |
| | | Blich-gyrB-R | AGAAACTTTTCNAGCGCTT | |
| | | RC | AAGCGCTTGAAAAGTTTCT | |
| *Tritrichomonas fetus* | 18SrRNA | TF-forward_18S | GTAGGTGAACCTGCCGTTG | 330-360bp |
| | | TF-reverse_5.8S | TTCAGTTCAGCGGGTCTTC | |
| | | RC | GAAGACCCGCTGAACTGAA | |

>Bovreproseq_synthpos_B. lich_gyrB_Adk_Trich_18s
AACCCTCACTATACGGAAGTGACGGGAACGTCTTCTAGATTTTGAGCAACGGAGAAAGCCTGTGAATGATGTCAGGAATGGCTT
ATCAGGGTAGGTCCTTCGCGTATATGTTGGACTTGTCTCTTATGCTTAGAACACGCTAATGCATACCGACCAAATAACGGAAATT
TAGTCGGGGGCGTAGCCCGCGTTGATGCATCCGGGCGGATTTAGCCGAACTCAGATCAGGTTGTTGCGCACGACATTCTTTATC
CCGAGCTCCTTTGTAGGTCGCCGGCTACCAATGGTAAAGTCGTTTCCCGGAGTGAGAATGGGGCACAAGCAAGCCATGATGTC
GGCCAATCCACTCGACCATCTGAACACGAGTGATCGGCAGAGGACAGCATAACGTAACGATTGAAAGGGTCCTTCGTATCGTG
CACGGTGTGGGCGTCTACTCAACCCGACTCCGCTAAGAATATCCATGGTTCGTCCTGAGAAGTCGTGCAAGCGCCTTCAACCAC
CACCATCATCAACATGGGGATCGCTAGGCTCCTACCCATATGAAACGGGATATCAGTACCCAGGCGTGTGGATACTAGGGAATC
GGGACGGACACTACAAAAGCGCTTGAAAAGTTTCTGTTCAATGTACATGGCGGTGTCGGAGGGTAAAGGGACACAGGCTGACC
GAGAGAGGGCCCCTTCATCTGTCGACATACCTATCCCAACTACGTAAGTCATATGTCTTCGCGGATACCGAAGCAGAATATCGCT
GGTTGACCTTACTATGCTACCATTTCACCTAGCCTATGGGAGCACACAAGGGACGGCGGGGACGCCATAATCTAGAGTGTGGAT
AAAACACGGGTCCCCATGCAGGGTCGAGGTCGGTCGTTATACAACCCAAGCAGTGATTTGAAGGGTCTAAGTTTCCCTAAGCAT
CCATCCTCGTAGTCGAACTCATCAATAGCATTAGATTACGCAAAGCCATGGATCCCATCTCACTAAGAACCTGCGTTTCAATATG
AACTCACAGATGAGAGCCTCTGGGATTCCCTACAGATCGTTCGCTAGTGCCATTCACTGGTAAGTCGTGAGATAGCGTTGAAGC
CACCTAGTTCGGAGAAACATCGGAACTCAGCAGGCTCGAGTTTATGACGTGCATCAAACCCAGCCTTTACTCGATTTATGACGT
GCATCGTTGTGTAGGTGAACCTGCCGTTGCAGAAGGCTTTCGCCCCCATTAATCGAGGTAATTGTACGTGTGCTGCTTCTTGCGT
AAGGTTCCGTGGCTGAGTCCAAGGAATTACTAACGCGAGCAGCGGGGCTGCTACTTGGATCAGGACTTCTGATAGGCTGGCCA
CCAGTAGCGCAGTTGTATAAACTAAAAGCCTGCGCCAGGCCTTGCTCCATGAATTCAGCGCGACGAAGGACTTCCCTACTGTTA
TTAATTTAAGCGACGAAGGACTGTACAGATGTCTCGGAATTCGTGGGGGAGTTGGAACCGGGGTGCCCCAGTGCAACTTCTTAT
GGTGGCACAACACCAAGAAAAGAAGACCCGCTGAACTGAAAATTAATGTCACTCGG

| Organism | Gene | Primer name | Primers (5'→3') | Size |
|---|---|---|---|---|
| *Trueperella pyogenes* | Pyolysin | Trueperella _pyogenes_ 493bp_ploNF | AACGGCCTTCTCGACGGTTG | 493 bp |
| | | Trueperella _pyogenes _493bp_ploNR | TAGCTCGGGTCTTGTTCAGG | |
| | | RC | CCTGAACAAGACCCGAGCTA | |
| | Tpyocpn60 | Cpn60-f3 | CGTTGAGGAGTCCAACAC | 280 bp |
| | | Cpn60-b3 | | |
| | | RC | GTCAACAAGATCCGTGGC | |
| *Chlamydia abortus* | Outer_membrane_protein | ch1-omp2 | ATGTCCAAACTCATCAGACGAG | 587 bp |
| | | ch2-omp2 | CCTTCTTTAAGAGGTTTTACCCA | |
| | | RC | TGGGTAAAACCTCTTAAAGAAGG | |

>Bovreproseq_synthpos_True_pyo_plo_cpn60_Chlamydia_omp2
TTGTCCCGACTCCCCTTCCAACGGCCTTCTCGACGGTTGGCACTCGATGAATGCCGGAGACGCCGGCCGCGTACGTAATATACC
GGCTCAGAGAACCGTAATTGTCAGGAGCGAGGGTCGGCACAACACTATGTGAACGCGGCCGATTTATCAGTGGTGTCCTCTTT
GACACGCCAGAACACATCGTCATTTTGAGTCATACCATCTTTGTCACTTTCGCAGACGAATTGCCATTATCTTCCAGTAATGAAAG
CATTACGTATGACCATCATAGTTACCTAGGGCGCCCATGAATGAAGTGCACGCTAAGTTGTCACTGGCGGAGGGGCTGCACACT
ATCATTCACCGTAGACGGACGTCGAAAATTCTCTGGTGATGTCGTGTATGCACTACAACAAATGGTGTCCTACTACATGTGATCT
TTGCCCGTTTGATGACACCTCAGAGGGTCTCGGCTTAGGCTCCTGCCAGTGTCAATCGTCATAGGCGCCTGAACAAGACCCGAG
CTACCAGGCTCGAGAGTCTAAGACGTTGAGGAGTCCAACACTTCTGTTCCGGAGACGCTGACAGAATCCTCGGTGGGTTCGCCT
GCTGCCGAAAGAATATTATTCGCTGATGCAGATCGCACGCCAGGGCATTACCGACAGGGAAGTAAGACACTCGATCGCTGTAG
GTCCCATACGAGTATAAAACTCTCGACGCTTACGCTAGGTTTAGACCGAGCGGTCAAGTGAAGCTTCATGAAGGTGGATCCGAT
CTCTATTGCGATGCCAATTTGCAGTAACCGCTAGGTCAACAAGATCCGTGGCTGCCTGCTAGGAATTCTAGGGATGTCCAAACTC
ATCAGACGAGTCTTGGACAACTTCGGGGGTGCCTCTGTTTCAAGGAATTAGTAGTATCTGAGGAAATAGGATATTGCGCCTCCA
TTGTAAGCATGGTCGGGGCCCACCTCGGCCTGAAAGCCCCCCAATCAACGGCCAAGTACGGACGCAGCCCACTGTGGACCCGT
GAGTGAACTCATCAACAGAAAGCCTGATCAATACACACGAAAACAAAGCCACGAAGTCTATTCGATGGCCAAAATCTGAGTGAC
TATTAGTCTTGGATCGGAGCCAGTAACTTGCCAGTGTGTTGACTCGCACTTACGGTGGTACATGCTTCCCGGCAGAAAGGCTCT
CCGCGACGTTTTTGGAACAGCTCATTGCCCTAAAATGCAACCCACCTTGTAGCTAAGTACCCAAACGACTGCGAATAACTCTCGA
CTATCTATCAGGGAGATGCCCGCAATGTTATACAGTTTGCGCAAGCGCTTACAAGCCCTTAATAGTGGAGGGTGAGAGGTTCCC
AGAAACTCGGAAATCCGCCTATGTGTTGGGATCGGAGACCCATCAGACTGGGTAAAACCTCTTAAAGAAGGCGATTTCGTCGCC
CGAGTCACTACT

| Organism | Gene | Primer name | Primers (5'→3') | Size |
|---|---|---|---|---|
| *Campylobacter jejuni* | Cj0414 | cj0414C1F | CAAATAAAGTTAGAGGTAGAATGT | 160 bp |
| | | cj0414-R | CCATAAGCACTAGCTAGCTGAT | |
| | | RC | ATCAGCTAGCTAGTGCTTATGG | |
| | hipO | HipO-F | GCTATAACTATCCGAAGAAGCCATCA | 350 bp |
| | | hipOR | GACTTCGTGCAGATATGGATGCTT | |
| | | RC | AAGCATCCATATCTGCACGAAGTC | |

>Bovreproseq_synthpos_Cjejuni_new
AATTCTATCGAATATTGACTCCACAAATAAAGTTAGAGGTAGAATGTGTCATGTACGCGAATTCTCTGGCTCAGCTTGTGTGAAT
TCGATATCCTAGGGAAAAGAGGCTCCGGCATGGGGCGGGAGCACGCCTTCAAGCCGGGACTGCTTATGGAATGGGAATCAGC
TAGCTAGTGCTTATGGGTAGAGCACGCATTACCAGAGTGCAGATTCATGTAGCTATAACTATCCGAAGAAGCCATCACGCCAGC
AATGAGCAGCCTCCGTATCCTCACACATTACTTAACAGTACTTTCAGAACCGCATTAGGTAAGGGGGTGGGCAAGTACCATGAC
CGACCAGAAGGGCACATGGTTACGAGTCCACCTACACAGGAGAGCTGGACGGAACTAAGGTTTTCACATAAGAATCGGGAGCA
AATAATCACTGTCTGTAACACAATCATCTGATCCGAATGTATTTGGAACTTATTGAGCATAAGTGCGCCGACAGATAAGCATTTC
GTGCCCTCCGTTTAATACGAGCGGCTTCTCCTAAAAAGCATCCATATCTGCACGAAGTCTATTACGACCGACAGACCTCTCTGCC
ACGGTCTCTGCCACGGTCTCGC

| Organism | Gene | Primer name | Primers (5'→3') | Size |
|---|---|---|---|---|
| *Trueperella pyogenes* | superoxide dismutase | sodA F | CGAGCTCGCCGACGCTATTGCT | 160 bp |
| | | sodA R | GAGCATGAGAATCGGGTAAGTGCCA | |
| | | RC | TGGCACTTACCCGATTCTCATGCTC | |

>Bovreproseq_synthpos_Trueperella_pyo_sodA_new
CAGCTAGTTCTTACAAGACCCTTCGAGCTCGCCGACGCTATTGCTGGTTCTCCGCGACTGCGTTTGCTCGTCACCAACATACAGG
CTACCAACCTCCCGAAAAAGCTGCACTTGCCACAGTAGCTTTAGAGTACCGCTATTTTACGCGAACTAGTCATACCGTGACTCCT
AAAGTCCAGATTGGACACAGAATGCAGTGGCACTTACCCGATTCTCATGCTCAATGCAGGTCGTCCCGCCGACATGCTACTGC

| Organism | Gene | Primer name | Primers (5'→3') | Size |
|---|---|---|---|---|
| *Mycoplasma bovis* | PPSM5-1 | PpSM5-1 | CCAGCTCACCCTTATACATGAGCGC | 442 bp |
| | | PpSM5-2 | TGACTCACCATTTAGACCGACTATTTCAC | |
| | | RC | GTGAAATAGTCGGTCTAAATGGTGAGTCA | |
| | uvRC | uvrC_MB-F3 | CCTGTCGGAGTTGCAATTGTT | 368 bp |
| | | uvrC_MB-B3 | CGGTCAACTTCAACTTGAATTTG | |
| | | RC | CAAATTCAAGTTGAAGTTGACCG | |

>Bovreproseq_synthpos_Mbovis_new
CCGACAGACCTCTCTGCCACGGCCAGCTCACCCTTATACATGAGCGCTCTCGCGCATTACGATACCCACTGTTAACTCGAAAGCA
CCGTCATCAGGCCTGATGTGAGAGATGGGAGTTCTGCATTAGTTTGTATTCACCTAACCCTGGGCTGGCTTTGCATAGCACTCTC
CTGCGCCGGCGACCAGATCCTGGTGATTTTTGACCCGCATACATATCGATACAAGTCCTACAGTTTGTTAGGGTGGCACATCGG
ACCTGCTACTGGGAGCCCCATGTATGATAGGGAATAGTAGAAGTCGGGATACTACCGGTGGGTGTTCGATGGCGAGTAGGCG
ACTCGGTAAACGTCTTGACGATCCCTTCTTATTTGTGCAATGGAAGGATTCCCTTTGCATTAACCTTCTTATTCGTCCAGCGAGAG
CACGTCGCTCCACGTGAAATAGTCGGTCTAAATGGTGAGTCAAGATCGGGCGAGGTTGTTTGACTCTAAAAGTATCAGTTCTAC
CCACTATACCTGTCGGAGTTGCAATTGTTGGATAATAAGGCGTTGAAGAGACTTAAAGCAGGATTTCTGGTTGTCTCACTTTCCA
CCCTCCTGCCGGCCGTCGTCTCTCGAGTTTTTGTTTAGGGAGCTGCCCTGTACGAAGTAAACACCGTGCCCATGTTCACTTGGAG
GCCCACGACACTAGCGTCAACGTATCCCTCTGGTTGGACTTTGTGAGCAGGGTGGTTGTACGTGCGCGAGTTGAGTTCTCAATG

TACCTGTGACAGTGGCCCCTCTGGCCAGCTTGCATTGCGGAAGTGCAGAGCGGAACGGAGCAAGTAAATGTCAAGTGCACCGA
CCTTATGTAAGTGAGTCAAATTCAAGTTGAAGTTGACCGGCGATGGTATTGACCTTGTAATGTATCACA
